# Separating direct, indirect and parent-of-origin genetic effects in the human population

**DOI:** 10.1101/2025.04.28.650988

**Authors:** Ilse Krätschmer, Laura Hegemann, Robin Hofmeister, Elizabeth C. Corfield, Mahdi Mahmoudi, Olivier Delaneau, Ole A. Andreassen, Archie Campbell, Caroline Hayward, Estonian Biobank Research Team, Riccardo E. Marioni, Eivind Ystrom, Alexandra Havdahl, Matthew R. Robinson

## Abstract

Here, we present a novel approach to estimate the degree to which the phenotypic effect of a DNA locus is attributable to four components: alleles in the child (direct genetic effects), alleles in the mother and the father (indirect genetic effects), or is dependent upon the parent from which it is inherited (parent-of-origin, PofO effects). Applying our model, JODIE, to 30,000 child-mother-father trios with phased DNA information from the Estonian Biobank (EstBB) and the Norwegian Mother, Father, Child Cohort (MoBa), we jointly estimate the phenotypic variance attributable to these four effects unbiased of assortative mating (AM) for height, body mass index (BMI) and childhood educational test score (EA). For all three traits, direct effects make the largest contribution to the genetic effect variance. But we find that parental indirect genetic effects make an equivalent combined contribution, and that there is a non-zero PofO effect variance for all traits. We calculate the heritability that would be obtained at the population-level in the absence of AM for common DNA loci, and show that the proportional contribution of direct effects to these heritability values can be calculated as 64.0% for EA in MoBa, 77.1% and 63.4% for height in MoBa and EstBB, and 81.2% and 88.0% for BMI in MoBa and EstBB. Additionally, using within-family genome-wide association testing, we identify 276 independently associated DNA regions that replicate across two additional biobanks, which all show a genotype-phenotype relationship that reflects an interplay of direct, indirect and PofO effects. Determining how direct, parental and PofO genetic effects combine across loci genome-wide to influence human phenotypic variation requires joint modeling of parental and child genotypes alongside the parental origin of loci and here, we make the first attempt to do this in the human population.

---

In many organisms, parental influence can be wide-ranging: from the environment created by parents for their offspring to develop within, to epigenetic DNA modifications that occur in parents during egg or sperm formation [1, 2]. In humans, these early-life parental effects persist in their phenotypic influence long beyond childhood, shaping adult health and disease [3].

Here, we focus on jointly estimating the degree to which the phenotypic effect of a DNA locus is attributable to alleles in the child (direct genetic effects), alleles in the mother and father (indirect genetic effects), or dependent upon the parent from which it is inherited (parent-of-origin effects, Figure 1). Indirect genetic effects arise because families differ in a wide range of characteristics that shape the environment in which a child develops. If these characteristics are correlated with parental genotypes, then the genotypes of both the mother and father will influence the traits of their children, beyond simply the alleles that are inherited. Where one copy of a gene in an individual (either from their mother or their father) is expressed to a greater degree than the other copy, genomic imprinting may also occur. This gives rise to parent-of-origin (PofO) effects, where the direct genetic effect of an allele at a locus is dependent on the parent from which it is inherited [1, 4]. As we describe below, the degree to which direct, maternal, paternal, and PofO genetic effects combine across loci genome-wide to influence human phenotypic variation has yet to be fully quantified.

**Fig. 1.**
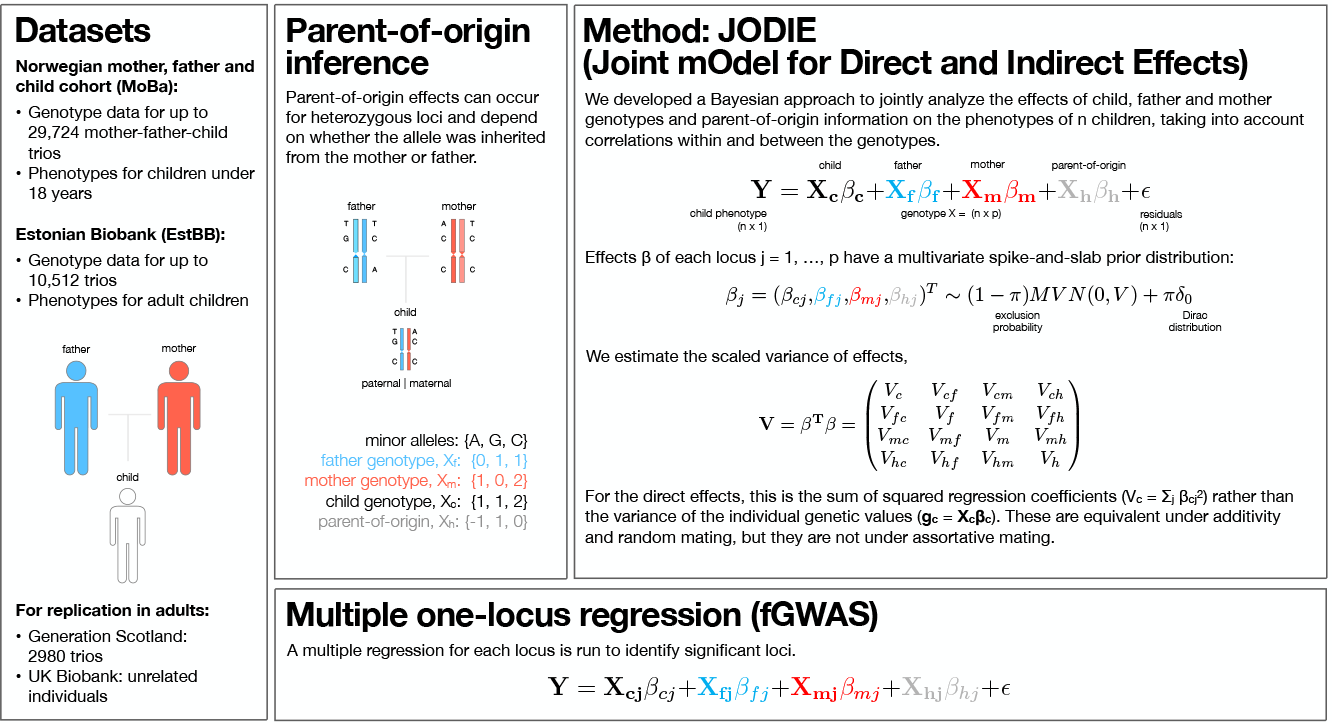
Jointly estimating direct, indirect maternal and paternal and parent-of-origin genetic effects, determining their contribution to phenotypic variance and identifying significant loci: used datasets, parent-of-origin inference, Bayesian method to estimate the scaled covariance matrix of effects and multiple regression method to identify significant loci.

## Results

We propose an iterative Bayesian variable selection regression model, that we call JODIE (Joint mOdel for Direct and Indirect Effects, see Figure 1 and Methods for a full description). The response variable is a trait observed in the child and the design matrix of the model is composed of the maternal, paternal and child genotypes of each trio alongside indicators for the parent-of-origin of each SNP inherited by reciprocal heterozygotes (1 for paternal inheritance, -1 for maternal inheritance, 0 for all other genotypes). The parental origin of heterozygous loci is determined by phasing the maternal, paternal and child genotypes of the trios. For each SNP, the model jointly estimates four regression coefficients for the child’s genotype (direct effect, ***β***_***c***_), the mother’s genotype (indirect maternal effect, ***β***_***m***_), the father’s genotype (indirect paternal effect, ***β***_***f***_), and the parent-of-origin indicator (parent-of-origin effect ***β***_***h***_). As all SNPs are fit jointly, the four regression coefficients of each SNPs are estimated conditionally on all other SNPs (see Methods).

As parents and children share DNA, estimating the direct effects of a child’s DNA requires conditioning on the variants in the parents. Since parents are the source of genetic variation in children, jointly modeling their genotypes alongside those of the child is sufficient to account for indirect genetically correlated variation. Including a vector of indicators of the parent from which the allele is inherited and modeling it alongside both the child and parental genotypes, is the only way to unbiasedly estimate PofO effects. This is because not every child genotype can be produced by all parental genotypes and thus conditioning on the variants in the parents is required to remove the correlation of PofO effects with indirect genetic effects [5]. Thus, we extend both previous PofO studies, which have not conditioned on parental genotypes [6, 7] and indirect genetic effect studies, which have not jointly considered PofO effects [8–13].

JODIE is a novel joint random effects modeling framework, where within each iteration, the SNP loci that contribute to phenotypic variance are identified and the four regression coefficients for standardized allelic variation in the child (direct, *c*), father (paternal, *f*), mother (maternal, *m*), and of the PofO heterozygote assignment (*h*) are estimated conditional on each other. From these regression coefficients, JODIE estimates the variance-covariance matrix, **V**, in each iteration of the algorithm as

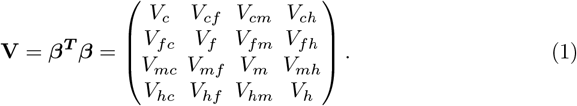

The diagonal of **V** is determined by the sum of the squared regression coefficients for each of the four components (***β***_***c***_, ***β***_***f***_, ***β***_***m***_, ***β***_***h***_), estimated genome-wide at each iteration. These values quantify the genic variance of each of these four components to phenotype, which is a scaled estimate of the variance of the effect sizes. As we show below, the values can be interpreted as the genic variance in a randomly mating (base) population. The estimated covariance (correlation) is a measure of the degree to which the four effects localize to the same regions of the genome.

We note that this differs from standard estimates of the SNP-heritability, 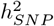, defined as the proportion of phenotypic variance attributable to the genetic markers. We can give an approximation of 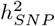 as 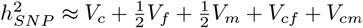 [11, 12, 14]. In what follows, when we discuss the variance attributable to direct effects, we refer to *V*_*c*_, likewise for *V*_*f*_, *V*_*m*_ and *V*_*h*_. When we mention the correlation of direct and maternal effects, we refer to the normalized values of *V*_*cm*_, likewise for all the other covariance terms.

We first show in extensive simulation studies, presented in the Supplementary Material, that the posterior estimates of ***V*** obtained by JODIE are unbiased, well-calibrated, and have the correct posterior coverage. Through the framework of simulation-based calibration and posterior predictive checking [15, 16], we assess the performance of JODIE across a range of generated child-mother-father trio sample sizes in Figures S1-S3. For samples sizes ranging from 10,000 to 30,000 trios, we show in Figure S1 that the posterior mean for all components of ***V*** returned by JODIE are unbiased estimates of the true simulated values. Figure S2 demonstrates that the standard deviation of the posterior distribution of each component of ***V*** estimated by JODIE conforms or is larger than the Poissonian error 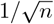. The Poissonian error directly reflects the uncertainty due to the sample size, while the estimated standard deviation is given by the square root of variation in posterior estimates across posterior iterations, thus only indirectly reflecting the sample size. We then show in Figure S3 that the posterior distribution returned by JODIE within each simulation replicate conforms to a uniform distribution. If the rank of the estimated 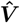 as compared to the true simulated value is uniform among the iterative posterior draws, 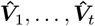, with *t* the number of iterations, then the sampler is well calibrated and the posterior intervals have appropriate coverage. This means that for any 90% posterior interval selected, the probability the true value falls in it will also be 90%. We show that JODIE provides a posterior with these properties over a range of trio sample sizes.

Finally, if a model captures the data well, then simulating new replicated datasets based on the fitted model parameters should yield a distribution with the same statistics as the original simulated dataset. To test the most critical part of our framework, the sampling of ***V***, we generate effects based on the ***V*** estimated by JODIE (input) and use these effects to resample ***V*** (output). The input and output distributions and their standard deviations agree well with each other for all components of 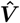, as shown in Figure S4, showing that the posterior distribution of 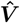 has appropriate coverage.

Having shown that JODIE performs as expected within generated trio data, we then expanded the simulation study to assess performance across: *(i)* five different scenarios for the values of ***V***; *(ii)* with and without assortative mating, and *(iii)* when simulating on top of real observed trio genotype data. As we wished to cover this wide range of scenarios and avoid unnecessary wastage of compute resources, we first show that the conclusions gained from our simulation study are identical with 10 replicates in each scenario as they are with 50 simulation replicates in Figures S5-S7. To aid interpretation in scenarios where the true components of ***V*** = 0, we propose a simple test for the significance of each variance component by calculating whether ≥ 95% of the posterior distribution of the sum of squared regression coefficients is ≥ 2%, known as a region of practical equivalence (ROPE) rule. We show in Figure S8 across 5 simulation scenarios that JODIE detects non-zero variance components and non-zero covariances with 100% accuracy under our ROPE rule and returns unbiased posterior mean estimates.

We explore the ability of JODIE to estimate ***V*** under both random and assortative mating as compared to other methods proposed in the literature. Through forwardin-time simulation across 5 generations, we first generate a phenotype under random mating where variation is attributable to only loci in the child (see Methods). In this setting, the sum of the centered and scaled child genotypes multiplied by their effects, Var(***g***_***k***_) = Var(***X***_***k***_***β***_***k***_), where k refers to child, mother, father or PofO, is equal to the true genic variance, as shown in the left top panel of Figures 2 and S9. This is because under random mating trait-increasing loci are, on average, independent. As a result, estimates obtained from Haseman-Elston (HE) regression, or Restricted Maximum Likelihood (REML) models fitting all four components together, and TrioGCTA [17], where direct and indirect parental effects are fit together with their correlations, are equivalent to the values obtained by JODIE, with all methods producing unbiased estimation.

**Fig. 2.**
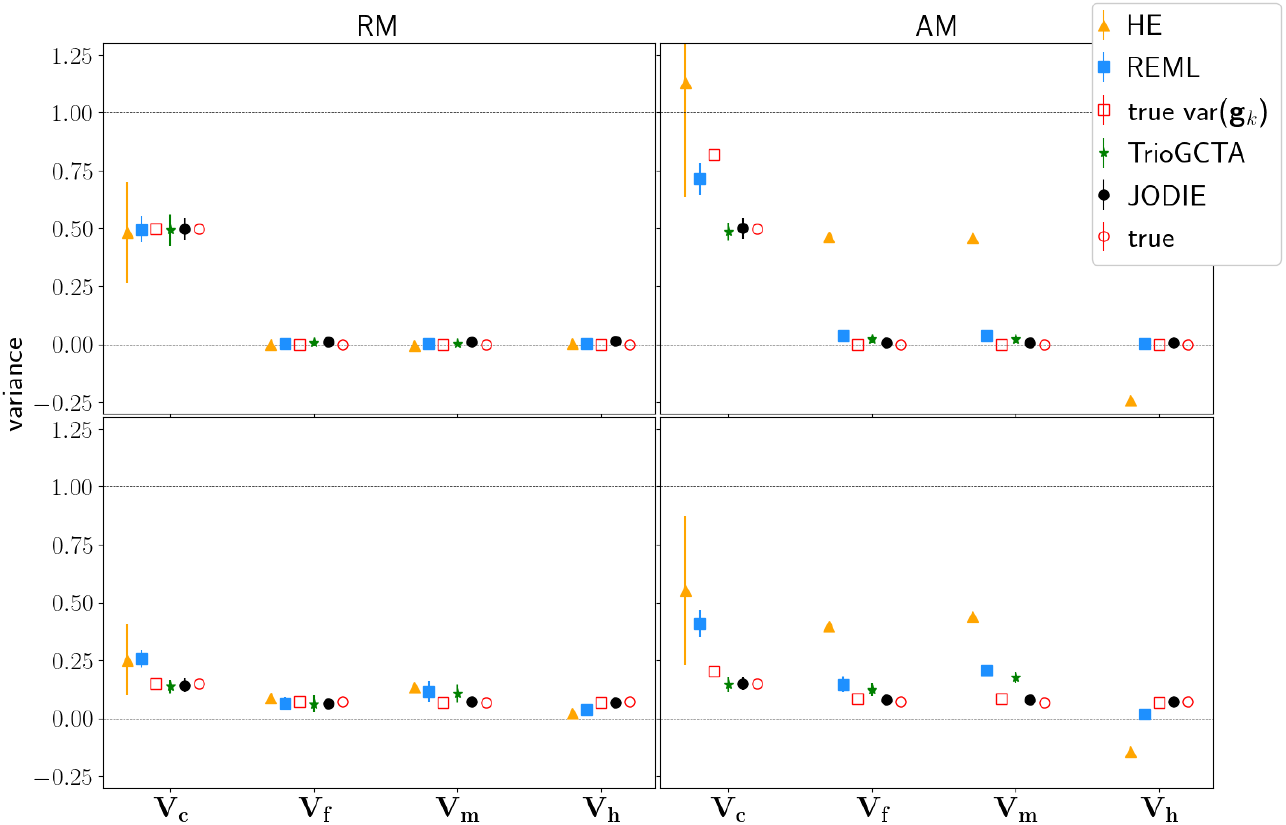
Simulation study of the influence of random and assortative mating on parameter estimation across statistical methods. For two different simulation scenarios, simulated true variances of direct, indirect parental and parent-of-origin genetic effects (labeled ”true”), true variances of the individual genotypic values multiplied by the effects, ***Xβ***, for the *k*^*th*^ component (labeled ”true var(**g**_*k*_)”) and the estimated variance parameters from Haseman-Elston regression (HE), restricted maximum likelihood (REML), trioGCTA (without PofO) and the JODIE model are shown. We generate genotype data, ***X***, of 16,000 child-mother-father trios for 52,310 common genetic variants randomly selected from 1000 Human Genomes data. Parents are mated randomly (random mating: RM), or are paired based upon their phenotypic values generating assortative mating (AM) with phenotypic correlation 0.5 across 5 generations. In the top panels, phenotypic data of each generation are generated where variation is only attributable to DNA loci in the child with direct effects. In the bottom panels, phenotypic data of each generation are generated where variation is spread across all genetic effects and correlated. The mean estimated variance across 10 simulation replicates is shown. Error bars indicate 2*×*standard deviation of the estimates across 10 simulation replicates.

Under different ***V*** scenarios under random mating, where there are parental indirect genetic and PofO genetic effects and their correlations (shown in the left bottom panel of Figures 2 and S9), we find that estimates using REML and HE regression become biased, over-estimating the proportion of variance attributable to direct genetic effects and underestimating the variance attributable to PofO effects, which is likely due to the highly correlated nature of the relationship matrices fit within the models and the finite sample size. TrioGCTA accounts for correlations within trios, but currently does not allow for PofO effects, which most likely results in the slight overestimation of one of the parental indirect effects. In contrast, JODIE estimates the genic variance of each component in an unbiased manner, controlling for the sharing of DNA among relatives.

Assortative mating (AM) can complicate inference as it increases the variance of the genetic values relative to that observed in a random mating population, by inducing positive long-range correlations among trait-increasing alleles genome-wide. Recent studies have focused on estimating: *(i)* the degree to which the variance of the genetic values is inflated under AM [11, 12, 18]; *(ii)* the contribution to variance of the average effects of untransmitted alleles in the parents [8–13]; *(iii)* the proportional contribution of indirect genetic effects to genome-wide association study (GWAS) estimates for a range of human complex traits [8, 11, 12, 17, 19] and *(iv)* PofO effects in large-scale data without accounting for parental indirect genetic effects [6, 7]. Here, our focus is different, as we jointly estimate the direct, indirect, and PofO effects, which is important because of the correlation of indirect and PofO effects. Note also that rather than estimating Var(***g***_***k***_), we estimate the variance-covariance matrix, **V**, as described above.

Under AM, the *k*^*th*^ diagonal element of **V** and Var(***g***_***k***_) for each of the four components are no longer equivalent. To demonstrate this, we conduct a simulation with only direct genetic effects where parents are paired based upon their phenotypic values generating AM with phenotypic correlation 0.5 across 5 generations (see Methods). We show that the equivalence of the true simulated variance of the genetic values, calculated as the sum of the centered and scaled child genotypes multiplied by their effects, Var(***g***_***c***_) = Var(***X***_***c***_***β***_***c***_), is no longer equal to the true scaled variances of the effects, 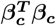, under AM, as can be seen from the right top panels of Figure 2 and S10. As a result, the estimates obtained from HE regression and REML are close to Var(***X***_***c***_***β***_***c***_). But the variance attributable to the maternal and paternal genetic effects is inflated and that attributable to PofO genetic effects is deflated in these settings, in particular for HE. When the phenotype is underlain by a more complex, correlated combination of direct, indirect and PofO effects, shown in the right bottom panel of Figures 2, S9 and S10, the inflation of true simulated variances of the genetic values, *V ar*(***g***_***k***_), is reduced, but the inflation, respective deflation, of the REML and HE regression estimates worsens and becomes severely biased. Correcting the HE and REML results for AM as suggested in Ref. [20] and shown in Figure S11 does not remove the bias due to AM in this case. TrioGCTA estimates the variance due to the direct effects correctly, but overestimates the variances due to the maternal and paternal genetic effects in all scenarios. In sharp contrast, under all scenarios, JODIE remains unbiased in the estimation of the scaled variance of effects, accounting for assortative mating.

Uniquely, JODIE is able to estimate covariances between all four genetic components. As shown in Figures S12 and S13, JODIE recovers the true value within the posterior distribution of the covariance estimate under RM and AM. Additionally, under our ROPE rule JODIE detects non-zero variance components (Figure S14) and non-zero covariances (Figure S15) with 100% accuracy, irrespective of whether mating is random or assortative.

In our final simulation study, we demonstrate that JODIE provides robust estimation in a realistic scenario where we simulate phenotypes from real genotype data of 10,512 trios from the Estonian Biobank. We show in Figures S16 and S17 that the posterior mean for all components of ***V*** returned by JODIE are unbiased estimates of the true simulated values. The exception is scenario “V4”, which is unrealistic as the simulated effects from each component are exactly uncorrelated, where the posterior mean diagonal elements (the sums of squared effects of the four components) are underestimates of the true simulated value by a few percent. This is likely due to the model favoring an estimate of zero when low-powered to resolve the effects across the different components. Nevertheless, Figure S18 shows that the posterior distribution returned by JODIE within each simulation replicate consistently conforms to a uniform distribution, showing that the posterior intervals of JODIE have appropriate coverage within this setting of real observed genotype data.

Taken together, we have simulated and analyzed 400 sets of large-scale trio data to ensure that the posterior estimates of 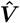 obtained by JODIE are unbiased, well-calibrated, and have the correct posterior coverage. Through the framework of simulation-based calibration and posterior predictive checking, we assess the performance of JODIE to be robust, especially in comparison to existing methods, across a range of sample sizes, different forms of ***V*** matrix, under random and assortative mating, and when simulating on top of both generated and real linkage disequilibrium patterns.

We then apply JODIE to ∼30, 000 child-mother-father trios with phased imputed genotype data in the Norwegian Mother, Father and Child cohort (MoBa) and ∼10, 000 trios in the Estonian Biobank (EstBB). For more details on the data, see Table S1. Figure 3 and Tables S2 and S3 show the estimated variances and correlations in MoBa and EstBB for height (HT), body mass index (BMI) and school educational achievement test scores (EA). Note that EstBB only holds data of adults, whereas MoBa contains phenotypes of children below the age of 18, from which we used ages 7-8 for HT and BMI and age 10 for EA. We find that indirect and PofO genetic effects have modest estimates of genic variance that are significantly different from zero. Maternal and paternal genetic effects make almost identical contributions to the phenotypic variance. Generally, direct genetic effects are stronger than either indirect parental or PofO effects, but the sum of the indirect parental effect and PofO variances is considerable in both populations.

**Fig. 3.**
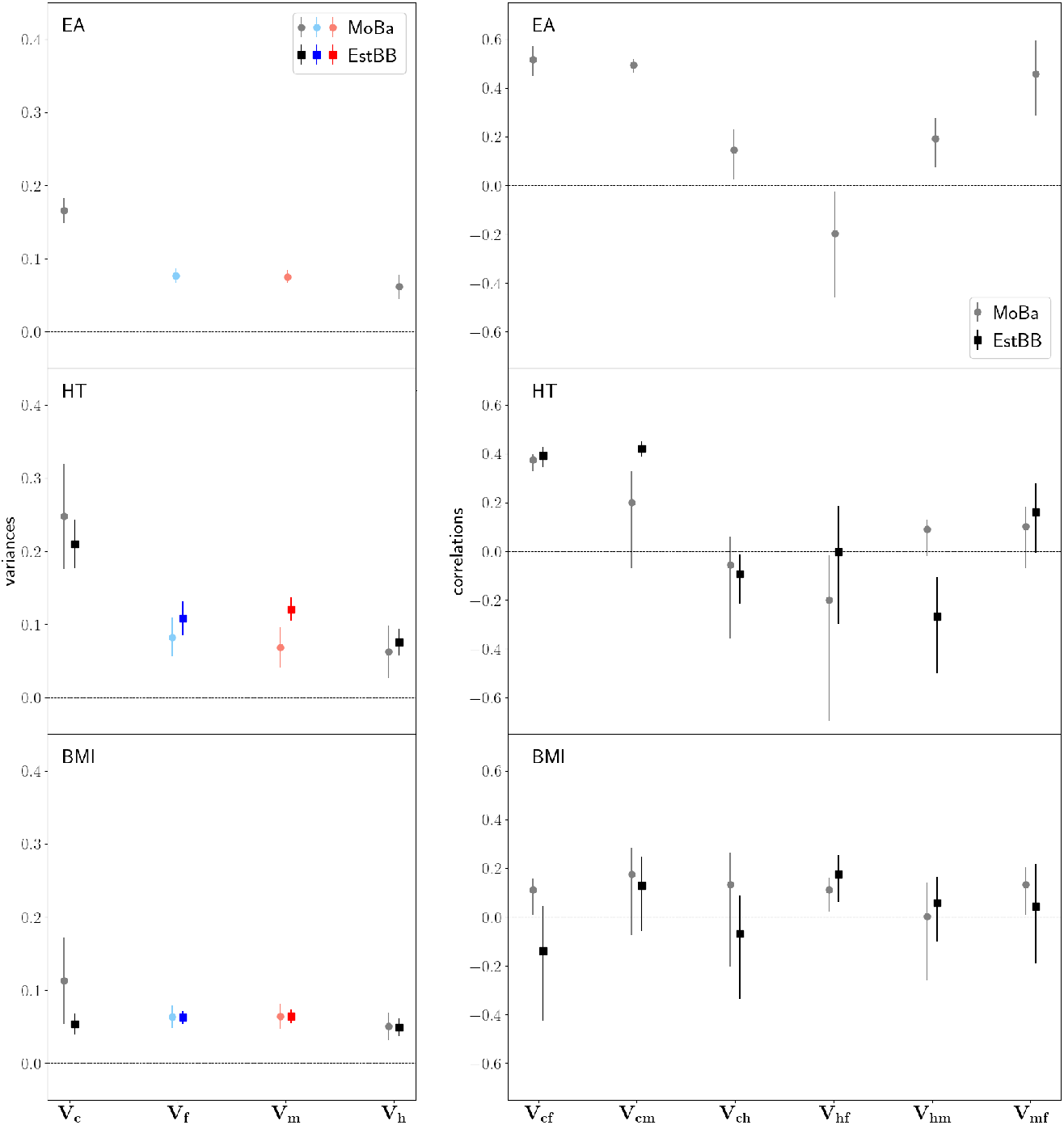
The contribution of direct, indirect parental and PofO effects across two biobank studies. Variances (left) and correlations (right) estimated by JODIE in MoBa and EstBB for education test scores at age 10 (EA, top), height (HT, middle) and body mass index (BMI, bottom). The left panels show the mean across 4000 posterior iterations (after burn-in) with uncertainties representing 95% confidence intervals. The correlations in the right panels are calculated as covariances scaled by the corresponding variances. The uncertainties are calculated using the posterior means ± 95% credible interval.

We find a positive correlation of the direct and maternal genetic effects, *V*_*cm*_, as well as a positive correlation of the direct and paternal genetic effects, *V*_*cf*_, across the genome for EA, height and potentially for BMI in MoBa for children aged 7-8 (Figure 3 right). This implies that similar loci underlie direct and indirect genetic effects within families. The correlation of the maternal and paternal genetic effects, *V*_*mf*_, is generally positive, but small. We note that this is not a measure of assortative mating for parental indirect effects, rather it is an estimate of the degree to which the same DNA regions underlie the maternal and paternal effects. All other correlations are not different to zero. Genome-wide, we find no consistent relationship between direct and PofO effects, *V*_*ch*_, across the three traits, which if they reflect imprinting, implies that the form of imprinting across the DNA is a combination of maternal, paternal and more complex patterns.

Generally, we find that estimates obtained from trios in MoBa for height and BMI are similar to those obtained from trios from EstBB, despite the fact that measures in MoBa are made on children and those in Estonia are made on adults. HE regression or REML applied to the same data, as shown in Figure S19, give far larger estimates of the variance attributable to direct genetic effects, which is consistent with what we show in our simulation study. Correcting the variance attributable to direct effects in EA and height for AM according to Ref. [20] using values for genetic similarities between partners in MoBa [21] only has a minor influence on the result.

We also conduct a within-trio one-marker-at-a-time genome-wide association study (family GWAS, fGWAS), where we jointly estimate the regression coefficients for allelic variation in the child, mother, father, and of the PofO heterozygote assignment. We run fGWAS in the MoBa and EstBB datasets separately and then meta-analyze the estimates (for more details, see Methods). Despite being underpowered to discover genetic variants given our sample size of up to ∼30, 000 trios for each trait, we find 11 independent (2MB distance and linkage disequilibrium R-squared ≤0.1) single nucleotide polymorphism (SNP) associations with EA, 15 independent associations with BMI and 276 independent associations with height, which span the genome and are not restricted to imprinted regions of the DNA (Supplementary Data Table 1).

The vast majority of these (9 for EA, 12 for BMI, 255 for height) replicate within 1MB distance and linkage disequilibrium R-squared ≥ 0.4 from a standard mixed-linear model association (MLMA) GWAS analysis within an independent sample, the UK Biobank, as shown in Figure 4a and Supplementary Data Table 1. We also find that almost all of the individual SNPs replicate in the Generation Scotland (GS) study for EA proxy phenotypes, height and BMI, within an fGWAS analysis of 2680 families. We show the distribution of the fGWAS test statistics in Figure 4b and Supplementary Data Table 1. Taken together, this confirms that the variants we find in fGWAS are associated with each of the respective traits (or a proxy related phenotypes) across four biobanks.

**Fig. 4.**
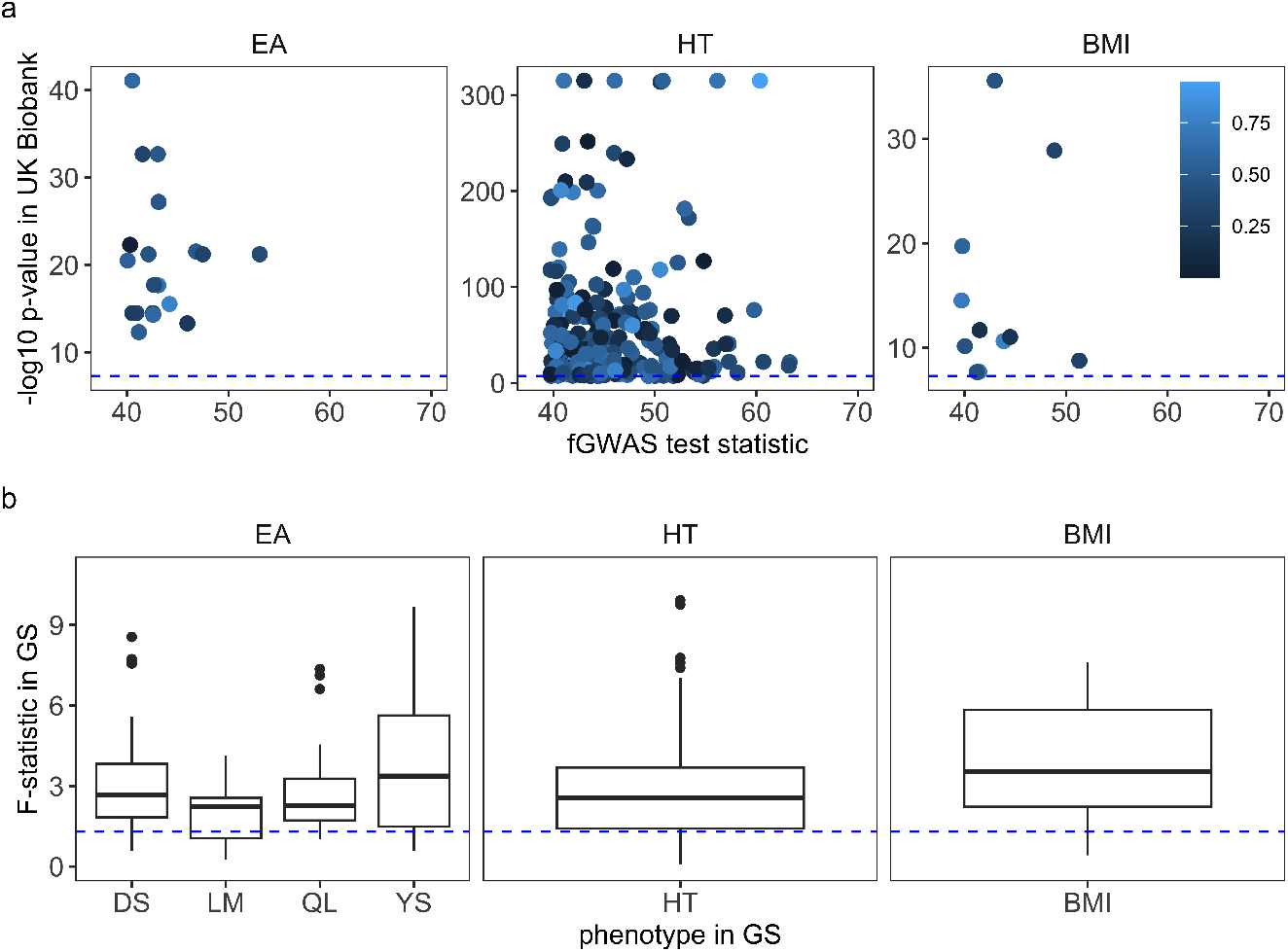
Replication of the family genome-wide association study (fGWAS) for education test scores (EA), height (HT), and body mass index (BMI) from the MoBa and EstBB studies, within the UK Biobank and the Generation Scotland (GS). (a) Genome-wide significant loci that are found in both MoBa and EstBB within the fGWAS analysis are tested to check whether they pass a significance threshold in a standard GWAS analysis (marginal mixed-linear model association analysis) in the UK Biobank. The −*log*_10_ of the p-value obtained for each SNP is shown on the y-axis. The x-axis gives the multivariate fGWAS test statistics. The point colors show the proportion of the z-score of the fGWAS direct genetic effect with respect to the sum of z-scores of all four genetic components. For the UK Biobank EA phenotype, we used a score of the maximum qualification achieved. Dashed blue line shows significance at the *p* = 5 × 10^−8^ level. We find that the majority of SNPs replicate as they are significantly trait-associated in three populations. (b) We use the SNPs that replicate in the UK Biobank to check if these SNPs also significantly explain variation in the trait in an fGWAS analysis of 2680 families in the Generation Scotland (GS) study. The fGWAS multiple regression gives an F-statistic of the variance explained by the regression for each SNP. The distribution of the test statistics is shown across SNPs, with dashed blue line showing significance at the 5% level. The EA trait in MoBa is approximated by digit symbol test (DS), logical memory test (LM), maximum qualification achieved (QL), and years of school education (YS) in GS.

To give an approximate indication of how much direct effects contribute to a certain locus compared to the other genetic components, we color the UK Biobank GWAS test statistic values according to the proportional contribution of the direct effects to the total effects in the fGWAS analysis in Figure 4a. This suggests that the 276 independent regions across the DNA that we identify in our fGWAS analysis are not only significantly trait-associated in UK Biobank because of direct effects, but rather that genotype-phenotype associations reflect a complex interplay of all genetic effects.

An alternative way to parameterize our one-marker-at-a-time model is to fit the effects of gametes rather than genotypes. In a gametic model, the transmitted and untransmitted alleles of the parents are fit jointly for each SNP. As we describe in the Methods, this has the advantage that transmitted and untransmitted alleles are uncorrelated (in contract to fitting the correlated genotypes of parents and children in the JODIE or fGWAS models), but the disadvantage that PofO effects have to be inferred from the estimated regression coefficients rather than being modeled directly. Figure 5 shows the parental and PofO regression coefficient estimated in the fGWAS analysis are correlated to those calculated in the gametic model, for loci that are genome-wide significant in MoBa, EstBB, UK Biobank and GS.

**Fig. 5.**
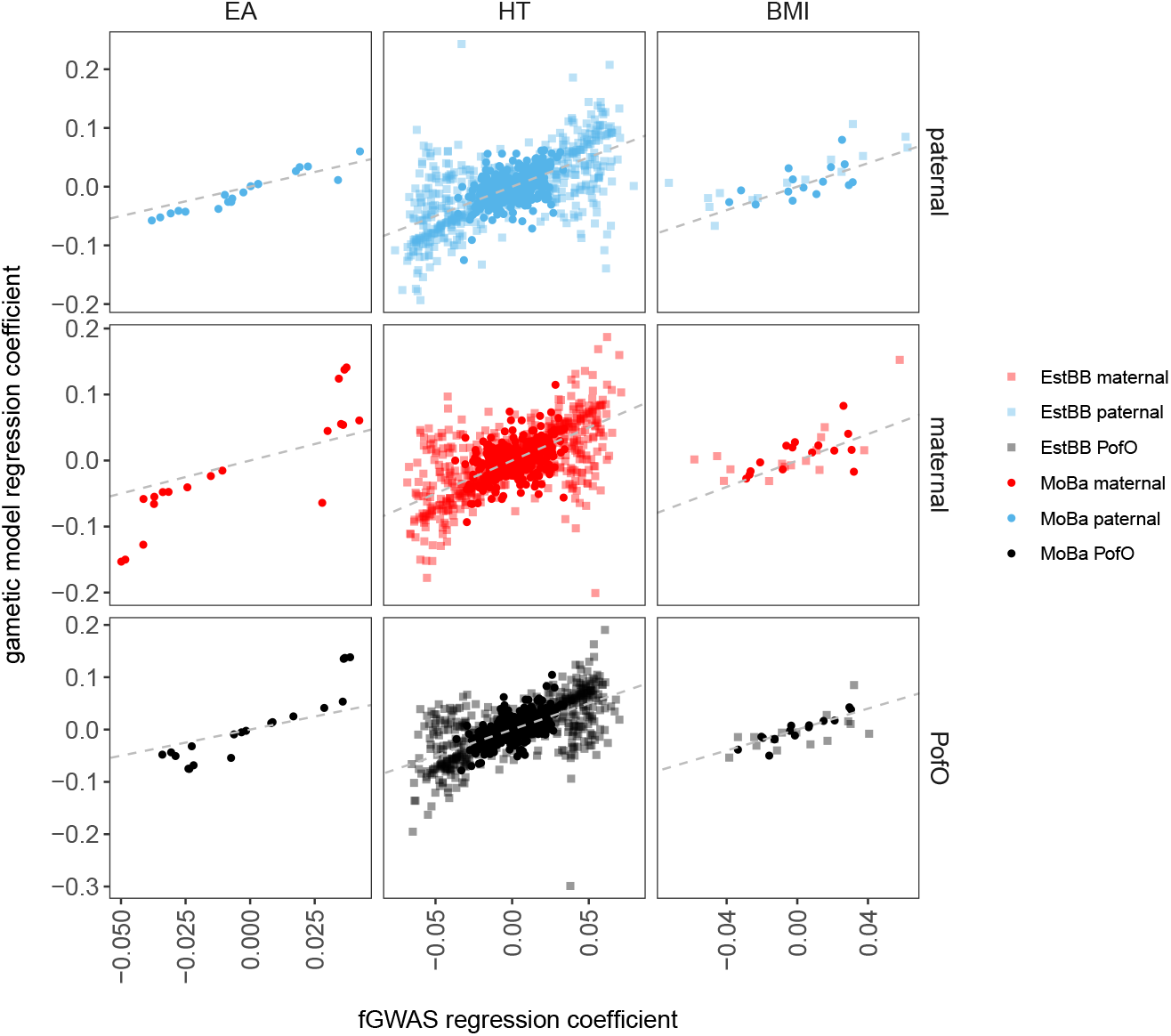
Family genome-wide association study (fGWAS) for education scores (EA), height (HT), and body mass index (BMI) from the MoBa and EstBB studies, estimated in an fGWAS and gametic model. For loci that are genomewide significant in MoBa, EstBB, the UK Biobank and GS, the regression coefficients of the paternal, maternal and PofO genetic effects are plotted when estimated using two different models. Estimates have a higher concordance, both from the 30,000 MoBa siblings (EA correlation = 0.949 paternal, 0.860 maternal, 0.915 PofO; HT correlation = 0.530 paternal, 0.517 maternal, 0.690 PofO; BMI correlation = 0.564 paternal, 0.553 maternal, 0.908 PofO) and from the 10,000 EstBB siblings (HT correlation = 0.648 paternal, 0.631 maternal, 0.658 PofO; BMI correlation = 0.818 paternal, 0.637 maternal, 0.604 PofO).

We highlight one locus per trait that are examples of the potentially complex nature of the phenotypic effects that are attributable to many DNA regions genome-wide. For each SNP, we show the relationships between the four genetic components shaping the phenotypic value of children in Figures S20, S21 and S22. The phenotypic means calculated for each child genotype change when parental genetic effects are taken into account, as is the case in both the JODIE and fGWAS models. In addition, the mean phenotypic values for heterozygous children are dependent upon whether they inherit the minor allele from their fathers or from their mothers.

We identify SNP rs13237829 as significantly associated with EA scores in MoBa. The SNP is near GTF2IRD1 (GTF2I Repeat Domain Containing 1) which has been shown to be associated with neuronal and synaptic development, craniofacial and cognitive development, and to underlie hypersociality symptoms in Williams syndrome [22, 23]. We do not have access to education attainment or cognitive measures within the EstBB, nor access to another cohort with comprehensive educational score measures from childhood. However, as well as replication in the UK Biobank for adult qualifications shown in Figure 4a (UK Biobank replication p-value 1.65e-06), we also replicate the effects of this SNP with 2680 adult families in the GS study, finding an association with years of school (GS replication p-value=0.004), and digit symbol (GS replication p-value=0.019) cognitive testing of adults (see Methods and Supplementary Data Table 1). This association is evidenced by the phenotypic mean values presented in Figure S20.

For height, we highlight SNP rs646806 discovered in MoBa and EstBB with p-value 1.11e-08 and replicated in the UK Biobank (p-value 1.49e-39) and GS (p-value 0.001). SNP rs646806 is located within the MGAT1 gene. MGAT1 encodes a transmembrane protein located in the medial compartment of the golgi apparatus. Mutations in these glycosylation proteins have been shown to have effects on the development of normal cells and dysregulation of several enzymes dependent on MGAT1 action are associated with human diseases [24]. MGAT1 is also associated with spermatogenesis and expressed in the testis [25], which suggests a link to paternal methylation imprinting occurring in spermatogenic cells. Figure S21 shows the phenotypic means in GS for child and parental genotypes for rs646806. Finally, for BMI, we highlight SNP rs919918 neighboring LINC01098. SNP rs919918 has MoBa and EstBB discovery p-value 2.14e08, UK Biobank replication p-value 4.29e-06, and GS replication p-value 0.003. This SNP is in LD with multiple SNPs (for example SNP rs2877501) in the GWAS catalogue within this region, which are listed as both BMI and insulin-like growth factor IGF-1 [26] associated. LINC01098 is also indicated in the Common Metabolic Diseases Genome Atlas as showing tissue-specific expression in the testis within GTeX data. Figure S22 shows the phenotypic means in GS for child and parental genotypes for rs919918. Taken together across traits, our findings highlight the potentially complex nature of the phenotypic effects that are attributable to many DNA regions genome-wide.

## Discussion

Here, we present a novel approach to determine the degree to which direct, maternal, paternal, and PofO genetic effects combine across loci genome-wide to influence human phenotypic variation. When partitioning the genome-wide genic variance into four components for EA, height and BMI, we generally find that direct genetic effects have the largest variance, and that the variances attributable to indirect parental genetic and PofO effects are not zero. The obtained variance patterns are consistent across the MoBa and EstBB biobanks. Uniquely, we can also estimate the covariances (correlations) between the genetic effects and genome-wide, we find that similar loci underlie direct and indirect genetic effects within families. Our one-SNP-at-a-time multiple regression (fGWAS) suggests that there may be hundreds of independent loci spread across the genome, where the genotype-phenotype association for EA, height, and BMI reflects a combination of direct, indirect and PofO effects. Almost all of the loci we identify also replicate across two additional biobanks, namely UKB and GS. We additionally confirm that parental and PofO genetic associations are similar when using an alternative gametic model parameterization. Taken together, our analysis of ∼30, 000 trios for each trait is the largest to date and provides significant insights into the genetic architecture of indirect and PofO effects.

Previous studies on indirect effects mostly focus on the degree to which population-level SNP heritability estimates are attenuated when adjusting for indirect genetic effects. From our results we can provide an approximation of the SNP heritability that would be obtained at the population-level in the absence of AM (the ”base” population), assuming that 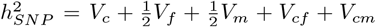. For the more than 1 million SNPs used in the JODIE model, we estimate 35.5% for EA in MoBa, 40.2% and 45.1% for HT in MoBa and EstBB, and 20.2% and 11.6% for BMI in MoBa and EstBB, respectively, which agree with previous findings [10, 27]. The PofO effects are not captured in standard SNP-heritability estimates, but their effects can be counted toward the direct contribution of a child’s DNA to its phenotype. Thus, the contribution of direct effects can be approximated as 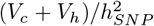, which is 64.0% for EA in MoBa, 77.1% and 63.4% for HT in MoBa and EstBB, and 81.2% and 88.0% for BMI in MoBa and EstBB. These calculations do not rely on missing/imputed parental genotypes, nor the use of the average of the parental indirect genetic effects. Rather, JODIE separates parental indirect genetic effects and estimates them alongside the direct and PofO effects, whilst uniquely modeling the covariances among the components, which are lacking from previous studies.

Our estimates of the contribution of direct effects are entirely in-line with those presented within previous studies for EA. For example, Kong *et al*. [8] estimates the proportion of direct effects over 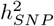 to be 65.8% for EA. Also, the genetic nurture effects and direct genetic effects on children’s educational attainment in eight cohorts estimated by Wang *et al*. [28] agree very well with our estimates for direct and indirect parental effects. The variance attributable to direct genetic effects for HT of 0.25 ± 0.07 is lower than previous estimates of 0.35 [11] and 0.34 [10], although the standard errors overlap our 95% credible intervals. Our estimates of the indirect effects are higher than those obtained from recent studies of indirect genetic effects for HT [12]. We suggest that this difference may be due to the fact that previous studies: (i) do not phase and impute data within-family; (ii) do not have both parental genotypes, so they frequently impute missing parental genotypes; and (iii) most studies use the average of the parental genotypes to control for average indirect genetic effects, which may all result in a reduction in the estimation of indirect effects. Indirect parental genetic effects for growth are found across almost all mammal species [1, 29]. The expectation in humans should be no different, with height being associated with economic status, health, and birth outcomes.

There are a number of limitations and remaining areas for improvement. First, the sample sizes of the biobanks we use, while representing the largest family-based cohorts in the world, are still very limited. Sample sizes of ∼30, 000 trios are insufficient to map large numbers of loci, or to produce genetic predictors with good accuracy for downstream inference tasks. Second, while our work represents an advance in variance component approaches for problems of this kind, we caution that with highly parameterized models and observational data, there may always be alternative interpretations for our empirical estimates, or many other models that could be applied which make different assumptions. Potentially, the effects estimated here may also broadly reflect any deviation from direct additive effects, including dominance (although dominance variance has been estimated to be small), heterogeneous variance (although we confirm parental and PofO effects remain under phenotypic transformation), subtle effects of LD, missing data (we have tried to minimize these using only full trios with no phasing error). Thus, it remains too early to conclusively determine the existence of and relationship among parental and PofO genetic effects, until large-scale repeated measures data are collected in early-life alongside where parental genotypes. Analyzing largescale data with JODIE would be hugely beneficial to understanding the genetic basis of complex traits in the human population, especially if data were collected across the range of human diversity.

One proposal in the literature is to phase large-scale extended families to obtain PofO information at scale [7]. However, studies in mice [1, 4], quantitative genetic theory and the results presented here, show that PofO effects and parental indirect genetic effects are correlated. Thus, any estimates of the PofO (or parental indirect genetic) effects without available parental genotype (or PofO) information will likely be biased. An additional key limitation is that we lack the data to model grandparental or wider family effects. Without conditioning on these, parental indirect genetic effects should not be interpreted as being causal in the parents. A final area for improvement is that our MCMC sampling scheme, which has acceptable computation times for current data sizes, could be replaced in larger-scale applications by variational inference approximations that would trade some accuracy for speed. Additionally, the model can be extended to utilize different prior formulations and sparsity over the components, to incorporate knowledge of minor allele frequency, linkage disquilibrium and genomic annotation, or to examine changes in genetic (co)variance through development within repeated measures data.

In conclusion, determining the degree to which direct, maternal, paternal, and PofO genetic effects combine across loci genome-wide to influence human phenotypic variation requires joint modeling of parental and child genotypes alongside the parental origin of loci. Here, we make the first attempt to do this in the human population.

## Methods

### The Norwegian Mother, Father, and Child cohort study

The Norwegian Mother, Father and Child Cohort Study (MoBa) is a population-based pregnancy cohort study conducted by the Norwegian Institute of Public Health. Participants were recruited from all over Norway from 1999-2008. The women consented to participation in 41% of the pregnancies. The cohort includes approximately 114,500 children, 95,200 mothers and 75,200 fathers. The establishment of MoBa and initial data collection was based on a license from the Norwegian Data Protection Agency and approval from The Regional Committees for Medical and Health Research Ethics. The MoBa cohort is currently regulated by the Norwegian Health Registry Act. The current study was approved by The Regional Committees for Medical and Health Research Ethics (2016/1702).

Information from the Medical Birth Registry of Norway (MBRN), a national health registry containing information about all births in Norway, and the MoBa questionnaire data were used to identify sex, year of birth, multiple births (in the offspring generation), and reported PO relationships. Prior to genetic analyses, all pedigrees were constructed based on the reported PO relationships for each pregnancy. Wherever possible, sex was assigned using information from the MBRN. In instances where sex was not specified in MBRN, the reported sex from the MoBa questionnaires was used.

Blood samples were collected from participating mothers and fathers at approximately the 17th week of pregnancy during the ultrasound examination. A second blood sample was taken from the mother soon after birth. The blood sample for the child was taken from the umbilical cord after birth. Biological samples were sent to the Norwegian Institute of Public Health where DNA was extracted by standard methods and stored. For more information on genotyping of the MoBa sample and for the family-based quality control pipeline used to prepare these data for analysis, see [30]. Here, we use the data generated after completion of module0 through module5 of the MoBa psychgen genotype imputation QC pipeline (see Code Availability) but we replace the QC and imputation pipeline used in Ref. [30], with our custom pipeline for phasing and imputation as described below.

The present study was conducted on a subset of the cohort of complete genotyped family trios (n = 29, 724) who who had data on at least one of the phenotype. Phenotypes used are national educational achievement test scores, height and body mass index (BMI). The national educational achievement test scores were ascertained via linkage to data from Statistics Norway. An average score across reading, English, and mathematics national tests taken at grade 5 (age 10-11) was used. Height and BMI were reported by mothers in the MoBa questionnaires. Height was collected in the 8year questionnaire. In case, no measure at the age of 8 was available, height reported at 7 years was used. If height was not reported at 8 or 7 years, height reported at age 3 years was used. Height at 8 years had a correlation at 0.85 with 7 and 0.70 with 3 years, respectively. BMI was collected in the 8-year questionnaire. For individuals without BMI reported at 8 years, BMI reported at 7 years was used. BMI at 8 and 7 years had a correlation of 0.70. For both height and BMI, values 3 standard deviations (SD) above and below the mean were removed. Values were standardized prior to combining across ages. All phenotypes were regressed on 20 genetic principal components, genotyping batch, child’s date of birth, age at data collection, and sex. The resulting measures were standardized to zero mean and unit variance prior to analysis.

### The Estonian Biobank

This project was granted ethics approval by the Estonian Committee on Bioethics and Human Research (https://genomics.ut.ee/en/content/estonian-Biobank). The activities of the Estonian Biobank are regulated by the Human Genes Research Act, which was adopted in 2000 specifically for the operations of the Estonian Biobank. Individual level data analysis in the Estonian Biobank was carried out under ethical approval 1.1-12/624 and 1.1-12/2856 from the Estonian Committee on Bioethics and Human Research (Estonian Ministry of Social Affairs), using data according to release application S16 from the Estonian Biobank.

As is common in many biobank studies, familial relationships were not known to us within the Estonian Biobank (EstBB). We calculated the coefficient of relatedness within the EstBB array data using the software KING (see Code availability). Parents were identified as individuals of opposite genetic sex who: (i) were unrelated (relatedness IBS0 < 0.0012), (ii) were within 10 years of age, (iii) both share half of their DNA with the same individual; and (iv) who are both at least 15 years older than the individual that they share half their DNA with. The latter two criteria automatically selected the children. We are confident that 10,512 true trio families were identified this way, which we used for our main analyses. Previous studies have also shown the accuracy of similar approaches to find trios [6, 31]. We additionally identified a further 26,209 putative mother-child duos and 6,364 putative father-child duos, by selecting pairs of individuals that share half of their DNA and who are at least 15 years apart. Within the sample were 2,373 sibling pairs. This gave 96,682 individuals in total, which were genotyped with Illumina Global Screening (GSA) arrays.

The Estonian Biobank individuals selected range from age 18 to 75 (mean age 30, 90% percentile age 44, 10% percentile age 20). Phenotypic values were selected from the baseline records for height and body mass index (BMI) and adjusted for genetic sex, age at measurement, and the first 10 principal components of the genetic data. We standardize all phenotypes to zero mean and unit variance prior to analysis.

### The Generation Scotland cohort

Generation Scotland (GS) is a large population-based, family-structured cohort of over 24,000 individuals aged 18–99 years. The study baseline took place between 2006 and 2011 and included detailed cognitive, physical, and health questionnaires, along with sample donation for genetic and biomarker data. Ethical approval for the GS cohort was received from the NHS Tayside Committee on Medical Research Ethics (REC Reference Number: 05/S1401/89) and Research Tissue Bank status was granted by the East of Scotland Research Ethics Service (REC Reference Number: 20/ES/0021). Participants provided written informed consent.

Genotyping array data are available for 20,026 (83%) of the original GS participants. Samples were genotyped using the Illumina HumanOmniExpressExome8V.1-2 A and HumanOmniExpressExome-8V.1 A arrays. Quality control measures were implemented, filtering out samples with a call rate of < 98% and SNPs with a call rate of < 98%, HWE of < 1 × 10^−6^ and MAF of ≤1%, leaving 20,026 samples and 630,207 SNPs.

We calculated the coefficient of relatedness within the GS array data using the software KING. Parents were identified as individuals of opposite genetic sex who: (i) were unrelated (relatedness < 0.0012), (ii) were within 10 years of age, (iii) both share half of their DNA with the same individual; and (iv) who are both at least 15 years older than the individual that they share half their DNA with. The latter two criteria automatically selected the children. We are confident that 2,680 true trio families were identified this way, which we used for our main analyses because we confirmed this using the available family information.

### UK Biobank

UK Biobank has approval from the North-West Multicenter Research Ethics Committee (MREC) to obtain and disseminate data and samples from the participants (https://www.ukbiobank.ac.uk/ethics/), and these ethical regulations cover the work in this study. Written informed consent was obtained from all participants.

We restrict our analysis to a sample of European-ancestry UK Biobank individuals to match the other cohorts analyzed. To infer ancestry, we use both self-reported ethnic background (UK Biobank field 21000-0), selecting coding 1, and genetic ethnicity (UK Biobank field 22006-0), selecting coding 1. We project the 488,377 genotyped participants onto the first two genotypic principal components (PC) calculated from 2,504 individuals of the 1,000 Genomes project. Using the obtained PC loadings, we then assign each participant to the closest 1,000 Genomes project population, selecting individuals with PC1 projection ≤ |4| and PC2 projection ≤ |3|. Samples were also excluded based on UK Biobank quality control procedures with individuals removed because of *(i)* extreme heterozygosity and missing genotype outliers; *(ii)* a genetically inferred gender that did not match the self-reported gender; *(iii)* putative sex chromosome aneuploidy; *(iv)* exclusion from kinship inference; *(v)* withdrawn consent.

We use genotype probabilities from version 3 of the imputed autosomal genotype data provided by the UK Biobank to hard-call the single nucleotide polymorphism (SNP) genotypes for variants with an imputation quality score above 0.3. The hardcall-threshold is 0.1, setting the genotypes with probability ≤0.9 as missing. From the good quality markers (with missingness less than 5% and *p*-value for the HardyWeinberg test larger than 10^−6^, as determined in the set of unrelated Europeans) we select those with rs identifier, in the set of European-ancestry participants, providing

a dataset of 23,609,048 SNPs. We select markers with MAF ≥ 0.002 to give 8,430,446 autosomal markers.

### Mendel phasing and haploid imputation

We phased the data using SHAPEIT5 [31] (see Code availability). We applied the option -pedigree of SHAPEIT5 to infer the parental origin of haplotypes for offspring alongside with phasing the data using Mendel phasing. For MoBa, we used only trios where parents and children were genotyped on the same genotyping array and conducted the phasing separately for each array in six sets. For EstBB and GS, as we wished to maximise both the number of trios and sibling pairs, we phased both trios and duos that were genotyped on the same array. Mendel phasing was thus conducted within-family and within-array, within each study.

We then matched the phased data to Genome Reference Consortium Human Build 37 (GRCh37) and swapped reference alleles to the Haplotype Reference Consortium data EGA release [32] (see Data availability). We then imputed SNPs in the parents and the children using IMPUTE5 v1.1.4 with the Haplotype Reference Consortium as a reference panel. As our data is phased, with each haplotype assigned to a specific parent, we used the parameter -out-ap-field to run a haploid imputation of the data and separately imputed the paternal haplotype and the maternal haplotype. As a result of haploid imputation, the PofO of imputed alleles can be probabilistically deduced from the imputation dosages: an allele imputed with a dosage of 0.9 on the paternal haplotype has 90% probability of being inherited from the father (i.e., PofO probability = 90%). This was done in 448 segments of the DNA for each set of data, and thus was also within-array. For each segment, we removed variants with an INFO score < 0.9, minor allele frequency < 1%, keeping G/T SNPs only after imputation.

Within each study, we concatenated files across segments for each chromosome. For MoBa, we then additionally merged the files across the six genotyping array sets for each chromosome. A final round of QC was conducted again, keeping only SNPs with missingness < 5%, INFO score > 0.9, minor allele frequency > 1% and HWE pvalue > 5x10^−8^. If there were multiple genotyping arrays within a biobank as in the case for MoBa, we calculated FST across the array sets and excluded SNPs that show any signs of being differentiated with FST value > 0.001. We also calculated the heterozygosity of each individual genome-wide using the plink –het command, and excluded individuals based on the F-statistics of heterozygosity.

After removing imputed SNPs with correlation *R*^2^ > 0.9 and MAF ≤ 0.005 in MoBa and ≤ 0.01 in EstBB, we converted the.vcf files to.zarr format, encoding the child, mother, father and parent-of-origin genotypic values. For children and their parents we used standard genotype scoring of 0, 1, and 2 counts of the minor allele. For the PofO effects, we scored reciprocal heterozygotes as 1 if the allele was inherited from the father (1 — 0), or -1 if the allele was inherited from the mother (0 — 1), with all other genotypes given a 0 value.

### Joint modeling of direct, indirect and imprinting effects

We developed a statistical framework (JODIE - Joint mOdel for Direct and Indirect Effects) that jointly estimates the direct (***β***_***c***_), indirect (***β***_***m***_ and ***β***_***f***_) and parent-of-origin (***β***_***h***_) effects on the child’s phenotype, as well as their variances and correlations in a way that controls for confounding of DNA sharing among relatives.

The standard statistical model in genetics considers *p* single nucleotide polymorphism (SNP) markers for *i* = 1, …, *n* individuals. The data is given in the form of an (*n* × *p*) matrix, **X**, where the elements are encoded as 0 for homozygous individuals at the major allele, 1 for heterozygotes and 2 for homozygotes at the minor allele. The relationship between the markers and the phenotype, **Y**, is assumed to be linear,

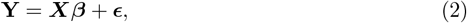

where **Y** and the residual error ***ϵ*** are vectors of length *n*. The effects ***β*** are given as a single vector of length *p*.

To take into account the parental and parent-of-origin effects on each individual, the genotypes and effects of mother and father as well as the parent-of-origin information are added to the model,

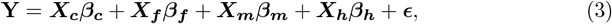

where the subscript c, m, f and h denote child, mother, father and parent-of-origin. The parent-of-origin information is encoded for the reciprocal heterozygotes as -1 if the minor allele is inherited from the mother and 1 if the minor allele is inherited from the father, all other genotypes are given a 0 value.

Equation 3 can be written in the same form as Equation 2,

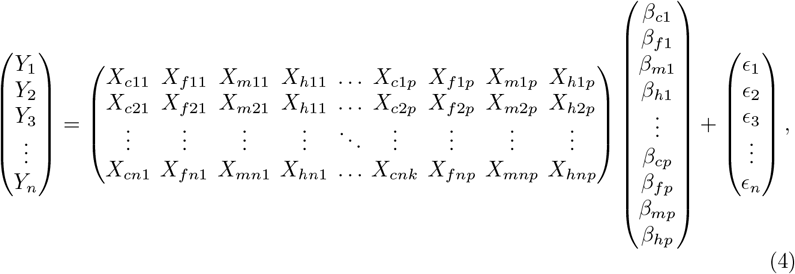

resulting in **X** = (*n* × *gp*) with columns standardized to have zero mean and unit variance, and ***β*** = (*gp* × 1) where *g* refers to the number of genetic components (*k* = 1, …, *g* and here *g* = 4). The phenotype vector **Y** is also standardized.

The effects in Equation 4 are estimated using a Gibbs sampler which is a Markov Chain Monte Carlo (MCMC) algorithm that is used when direct calculation is difficult. Since we want to take into account correlations between the SNP markers (linkage disequilibrium, LD), we adopt a Bayesian approach and place priors on the model parameters. The phenotypes and residuals are modelled with a multivariate normal distribution,

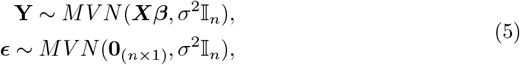

where *σ*^2^ denotes the variance, 𝕀_*n*_ is the identity matrix with dimensions (*n* × *n*), and **0**_(*n×*1)_ is a vector of zeros of length *n*.

A spike-and-slab prior is assumed for the direct, maternal, paternal and parent-of-origin effects of each marker *j*, ***β***_*j*_ = (*β*_*cj*_, *β*_*fj*_, *β*_*mj*_, *β*_*hj*_)^*T*^, i.e. they are assumed to be either distributed as a multivariate normal (MVN) or a Dirac delta, *δ*_0_,

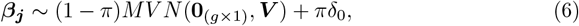

where the prior probability of marker, *j*, not being included in the model, *π*, is Dirichlet distributed and ***V*** is the genetic (co)variance matrix. The direct, maternal, paternal and parent-of-origin effects are estimated jointly, such that the effects at each marker are estimated conditionally on all other markers.

The covariance matrix is modelled as outlined in Section 2 of Ref. [33], using a modified Cholesky decomposition,

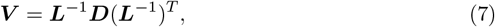

where ***D*** represents a diagonal matrix with only positive elements *d*_*k*_ (*k* = 1, …, *g*),

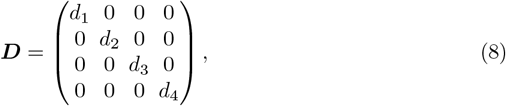

and ***L*** is a lower triangular matrix with diagonal elements set to 1,

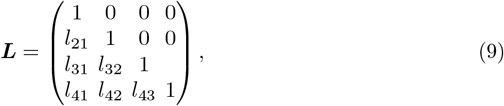

for *g* = 4. The diagonal elements of ***D*** are assumed to be inverse Gamma distributed,

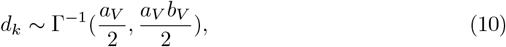

where *a*_*V*_ and *b*_*V*_ refer to the prior shape and scale parameters. The values of *a*_*V*_ and *b*_*V*_ are chosen to be *a*_*V*_ = 2 and *b*_*V*_ = 0.1. The elements *l*_*i*_ of each row *k* of ***L*** are modelled with a multivariate normal distribution,

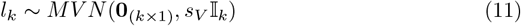

where the prior variance *s*_*V*_ is set to 0.001. This parametrization of the covariance matrix is preferable to the inverse Wishart distribution because off-diagonal elements in ***L*** are unrestricted.

The residual variance *σ*^2^ is modelled with an inverse Gamma distribution,

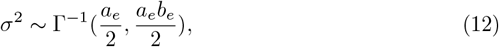

with prior scale and shape parameters, *a*_*e*_ = 1/*n* and *b*_*e*_ = 0.1.

Note that the formulas above are shown for four genetic components, as we are interested in the effects of direct, indirect paternal and maternal and PofO effects on phenotypes. However, JODIE is not limited to four components, but can take any number of genetic components above 1, *g* > 1.

The Gibbs sampler performs the following steps for each iteration:

1. Sample the intercept from a normal distribution.
2. Randomly pick a marker *j* and sample ***β***_***j***_ from its conditional posterior distribution

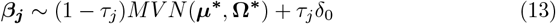

with posterior covariance

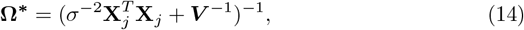

posterior mean

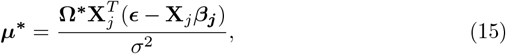

and posterior exclusion probability

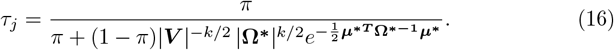
3. Repeat step (2) until all markers are sampled.
4. Sample the exclusion probability *π* from *Dirichlet*(*p* −*Z, Z*), where *p* is the total number of markers and *Z* is the number of non-zero markers.
5. Calculate ***V*** = ***L***^−1^***D***(***L***^−1^)^*T*^ by:
  a. Sampling the diagonal elements of ***D*** from

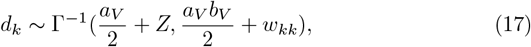

where *w*_*kk*_ is element (*k, k*) of ***w*** = *Z****Lβ***^*T*^ ***βL***^*T*^.
  b. Sampling the elements of the lower triangular matrix ***L*** from a multivariate normal distribution with mean

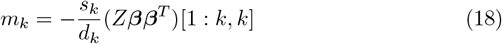

and variance

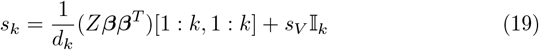

for each row *k* of ***L*** where [1 : *k*, 1 : *k*] denotes the submatrix of (*Z****ββ***^*T*^) between rows 1 to *k* and columns 1 to *k* and *s*_*V*_ is the initial variance of the multivariate normal.
6. Sample the residual variance from

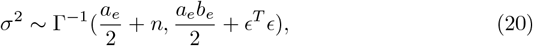

The number of iterations is set to 5000, including a burn-in period of 1000. Simulations as well as real-data studies show that the MCMC reaches convergence after only a few hundred iterations. The burn-in period of 1000 is chosen to be well away from this point to make sure that the values for posterior means are only taken in the region of convergence. Multiple chains are run after the burn-in period for the estimation of the posterior means. The uncertainties of the posterior means given throughout the paper represent the variation across the posterior iterations and thus only indirectly reflect the sample size.

The time complexity of JODIE is of order O(npg). The sampler is currently set up so that the full dataset needs to be read into memory. For further details on the implementation of the sampler using a Bulk synchronous parallel Gibbs sampling scheme with message passing interface [34], see links in Code availability.

We compare the estimates of JODIE in Estonian and Norwegian trios to HE regression and REML. We have chosen not to compare to trioGCTA as adding a parent-of-origin component in this approach is not feasible, we have RAM restrictions on our access to the EstBB data, and the error bars in the current R implementation seem unreliable in cases where the variance is close to 0 or 0.

### Modeling direct, indirect and imprinting effects of each SNP

The effect sizes and posterior inclusion probabilities estimated for each SNP by JODIE are conditional on all other SNPs genome-wide. We also determined conventional marginal one-SNP-at-a-time estimates, using the following within-trio GWAS model:

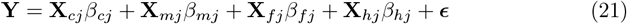

where for each SNP *j*, the direct and parent-of-origin effects are estimated conditional on the parental genotypes at that SNP. As the four regression coefficients *β*_*cj*_, *β*_*fj*_, *β*_*mj*_, *β*_*hj*_ are estimated jointly for each SNP *j*, we conducted joint significance testing using a multivariate *χ*^2^-test. For each SNP, we calculated a z-score for each effect by dividing the estimates by their standard error, SE:

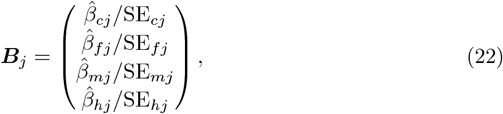

We then calculated 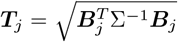, where **Σ** is an identity matrix. Under the null hypothesis that the z-scores of the four components for each SNP follow a multivariate normal distribution, where each has variance 1 and the correlation among components is 0, asymptotically 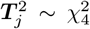, a Chi-squared distribution with *g* = 4 degrees of freedom. SNPs will pass the significance threshold when either one component has a sufficiently high z-statistic that passes the threshold, or where there are multiple components with sufficiently high z-statistic values that jointly pass the threshold. We also calculated ***T***_*j*_ where we use the regression coefficients directly without dividing by their SE for ***B***_*j*_ and where **Σ** is the empirical covariance matrix of the estimated regression coefficients across markers. Under the null hypothesis that the regression coefficients of the four components for each SNP follow a multivariate normal distribution, described by the covariance matrix **Σ**, asymptotically 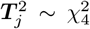, a Chi-squared distribution with *g* = 4 degrees of freedom. SNPs will pass the significance threshold when the components have regression coefficients that differ to the covariance observed across the DNA.

For height and BMI, we meta-analyze the estimates from the *s* = 2 datasets, EstBB and MoBa. We use a generalized least squares (GLS) method because the (co)variances of the effect estimates from the different studies are unequal. For each study and each SNP, we have a vector of effects:

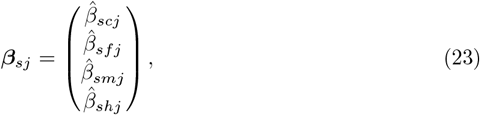

and a covariance matrix across SNPs for each study **Σ**_*s*_. We stack the *s* vectors ***β***_*sj*_ to get a long vector with length *sp*

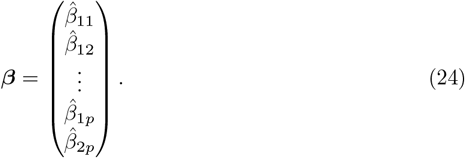

Assuming that the *s* studies are uncorrelated, we create a blockwise diagonal matrix for the covariance of the stacked vector ***β***

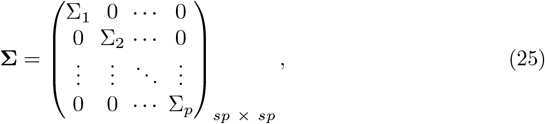

The fixed effect meta analysis estimator is then

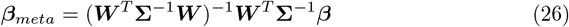

with ***W*** a stack of *s* identity matrices of size *g* × *g*. The standard error of the estimates is then

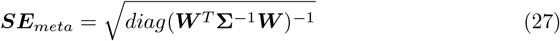

and the test statistic is 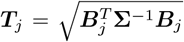, with **Σ** as *diag*(***W*** ^*T*^ **Σ**^−1^***W***)^−1^. The alternative is **Σ** = (***W*** ^*T*^ **Σ**^−1^***W***)^−1^ we present both within Supplementary Data Table

1. We checked the heterogeneity of the test statistic estimates with a test statistic

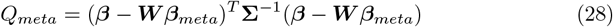

where under the null hypothesis of no heterogeneity the scalar *Q* asymptotically follows a Chi-squared distribution with *g* degrees of freedom.

### Replication within the UK Biobank and GS

In the UK Biobank, we select standing height (UK Biobank field 50), body mass index (UK Biobank field 21001) and a numerical ranking of education qualification (UK Biobank field 6138). Prior to analysis, all phenotypic values were adjusted by the participant’s recruitment age, their genetic sex, north-south and east-west location coordinates, measurement center, genotype batch, and the leading 20 genetic PCs and then scaled to zero mean and unit variance. We analyze the data using the mixed-linear model association testing framework of gVAMP [35] fitting all 8,430,466 SNPs in both the first and second steps, and check the test statistic values of the SNPs that were discovered in the fGWAS analysis of MoBa and EstBB. We also confirmed the test statistics values by running REGENIE [36] using a subset of 880k genetic variants in step 1 and all 8,430,466 SNPs in step 2. Here, we set a replication p-value threshold of 5 × 10^−6^.

In the Generation Scotland (GS) study, we analyze the following phenotypes: bodymass-index (BMI *kg/m*^2^); height (cm); logical memory (verbal declarative memory, LM), calculated from the Wechsler Logical Memory test by taking the sum of immediate and delayed recall of one oral story [37]; digit symbol (DS), ascertained from the Wechsler Digit Symbol Substitution test in which participants recoded digits to symbols over a 120 second period [37]; year spent in education (YS); and finally highest educational qualification achieved (QL). Years spent in education was self-reported as the total years attended school/study full-time, with coding 0: 0, 1: 1-4,2: 5-9, 3: 10-11, 4: 12-13, 5: 14-15, 6: 16-17, 7: 18-19, 8: 20-21, 9: 22-23, 10: more than 24 years. For highest educational qualification, participants were asked for the highest educational qualification they have obtained, with data then coded as: 1 - College or University degree, 2 - Other professional or technical qualification, 3 - NVQ or HND or HNC or equivalent, 4 - Higher Grade, A levels, AS levels or equivalent, 5 - Standard Grade, O levels, GCSEs or equivalent, 6 - CSEs or equivalent, 7 - School leavers certificate, 8 - Other, 9 - No Qualification. All phenotypic data were available for the children of all of the 2680 trios and values were standardized to mean zero and variance one prior to analysis. We ran the fGWAS model described above for the subset of SNPs that: *(i)* we identified as significantly associated with EA, HT, and BMI in the fGWAS analysis of MoBa and EstBB, and *ii* that replicate in the GWAS analysis of the UK Biobank. Here, due to the small sample size and the fact the we are conducting a replication analysis, we set the replication p-value threshold to 0.05.

### Alternative parametrization with the gametic model

Genotypes of parents and children are correlated. By splitting the genotype at each locus *j* in transmitted (***T***) and untransmitted alleles (***U***), we can ensure that the data is uncorrelated. However, this gametic model does not allow to explicitly model the PofO effects.

The gametic model is:

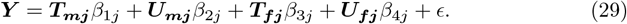

Linking the effects of the genotype model in Eq. 21 with the gametic model results in

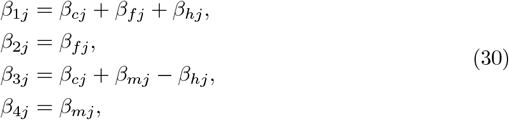

assuming that each parent has PofO effect *β*_*h*_ of the same size, but opposing direction on their child. Thus, we can directly confirm the indirect maternal and paternal effects of the gametic model. The PofO effect can be calculated as

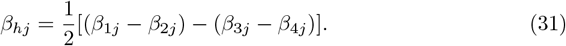

### Simulation study

A simple, but realistic forward in time simulation (based loosely on Ref. [38]) is used to created genotype and phenotype data for parents and children. We begin by simulated genotypes of unrelated individuals in generation 0 with realistic LD based on 52,310 randomly selected SNPs from the 1000 genomes project data (http://ftp.1000genomes.ebi.ac.uk/vol1/ftp/release/20130502/). Individuals are paired at random to produce two offspring each by recombining the parents’ haplotypes based on a genetic map (release b37, for example from https://github.com/odelaneau/shapeit5/tree/main/resources/maps/b37) and a single offspring is selected at random. Effects were generated for 1000 markers with a multivariate normal distribution with mean 0 and variance **V** scaled by the number of causal markers and randomly assigned for each simulation. Multiplying the simulated effects with their respective standardized columns of each **X** matrix (**X**_*c*_, **X**_*f*_, **X**_*m*_, **X**_*h*_) and summing the values, we obtained a genetic value, **g**, for each individual. In each scenario, a vector of residuals was sampled from a normal distribution with variance 1 − *var*(***g***), and added to **g** to obtain a vector of phenotypes, **Y**.

Using this simulation setup, we first examined whether the posterior estimates of ***V*** obtained by JODIE are unbiased, well-calibrated, and have the correct posterior coverage. Through the framework of simulation-based calibration and posterior predictive checking, we assess the performance of JODIE across a range of generated child-mother-father trio sample sizes ranging from 10,000 to 30,000 trios, using a realistic variance scenario, which is similar in pattern to what we estimate in EstBB and MoBa:

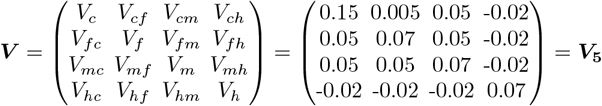

We create 50 replicate datasets for each of 10,000, 15,000, 20,000, 25,000, and 30,0000 individuals. For each of these 250 datasets, JODIE was run for 1500 iterations. The posterior mean estimates of all components of ***V*** were calculated using the last 1000 iterations.

To asses the soundness of JODIE as a posterior sampler, we check the difference between the posterior mean estimates of each element of 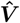 and the underlying simulated values and the standard deviation (SD) of the posterior of each element of 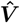. The standard deviation estimated by JODIE is the variation in the posterior sampling process of 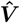 across posterior iterations. Its size is similar to the Poissonian error 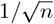, that directly reflects the uncertainty due to the sample size. Moreover, we use standard approaches of simulation-based calibration (SBC), a general procedure for validating inferences from Bayesian algorithms. If our estimated posterior 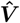 is distributed as a draw from the expected posterior, then the rank statistics of the true simulated ***V*** with respect to all of the posterior estimates 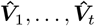, with *t* the number of iterations, should be uniform and if so, then it follows that posterior intervals will have appropriate coverage. Thus, for any 95% interval selected, the probability of the true ***V*** value falling within it will also be 95%.

A simple *χ*^2^ test for uniformity can be formulated by binning the ranks 1, …, *t* into *M* bins and testing that the bins all have roughly the expected number of draws in them. If *a*_*l*_ is the number of ranks that fall into bin *l* and *e*_*l*_ is the number of ranks expected to fall into bin *l*, which is proportional to its size under uniformity, then the test statistic is 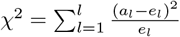 with *L* − 1 degrees of freedom.

We conduct further posterior predictive checks for 10 replicated datasets of 15,000 trios. Posterior predictive checks are a way of measuring whether a model captures relevant aspects of the data by simulating new replicated datasets based on the fitted model parameters and then comparing statistics applied to the replicated dataset with the same statistic applied to the original dataset. Here, we are interested in whether the minimum, maximum and SD of the posterior distribution of 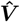 are a true reflection of the distribution of 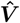 expected when sampling the variance-covariance matrix under the multivariate normal distribution.

We then compare simulation-based calibr samples sizes ranging ation, the distribution of the posterior mean estimates of each element of 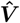 and SD of the posterior of each element of 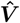 for different number of simulation replicates, ranging from 5 to 50. These studies show that each posterior distribution is well calibrated, and a sample of ten datasets is sufficient to demonstrate that this calibration is repeatable. Given this, we then created a further 10 datasets for a range of five 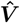 scenarios:

1. Scenario 1 represents a covariance matrix with only direct effects:

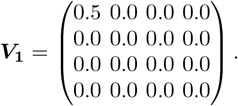
2. In the second scenario, direct as well as parent-of-origin effects are contributing to the variance:

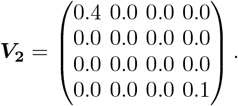
3. Scenario 3 assumes direct effects as well as smaller indirect maternal and paternal effects:

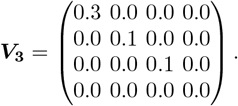
4. In scenario 4, the total variance of 0.5 is spread over all four genetic components. There are no correlations.

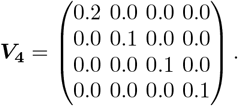
5. The last scenario represents a realistic data scenario, a similar pattern to what we have measured in EstBB and MoBa:

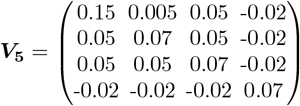

Effects and phenotypes for 16,000 trios are generated in the same way as described above. We run JODIE for each of the 50 simulated datasets for 1500 iterations. The posterior mean estimates of all genetic (co)variance components were calculated using the last 1000 iterations.

We then considered assortative mating by extending our simulation forward in time for 5 generations where the offspring of generation 0 become the parents for the next generation. Parents after generation 0 mate based on their ordered phenotypes to create assortative mating. The phenotypes for ordering are calculated as:

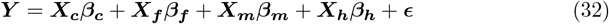

where *ϵ* is a value drawn from a Normal distribution with mean 0 and standard deviation of 0.01 to facilitate ordering. The phenotypes are ordered to introduce a correlation, *ρ* = 0.5, between the parents. This procedure is a simplified version of the unknown realistic process. For the implementation of the AM procedure, see Code availability. We simulate forward in time for ten replicates of each of the five ***V*** scenarios and apply HE regression, REML, trioGCTA and JODIE for each of the 50 datasets.

Finally, using the real observed SNP data of EstBB chromosome 2 (*p* = 89, 578 markers, n = 10,512 individuals), we simulate ten replicates of each of the five ***V*** scenarios. We compare the 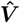 estimates obtained by JODIE to the truly simulated one for each of these 50 simulated datasets.

## Supplementary information

Supplementary Information accompanies this work.

## Acknowledgments

We thank Zoltan Kutalik, Peter Visscher and members of the Robinson group at ISTA for their comments, which improved this manuscript. This work was funded by an SNSF Eccellenza Grant to MRR (PCEGP3-181181), and by core funding from the Institute of Science and Technology Austria.

The Norwegian Mother, Father and Child Cohort Study is supported by the Norwegian Ministry of Health and Care Services and the Ministry of Education and Research. We are grateful to all the participating families in Norway who take part in this on-going cohort study. We thank the Norwegian Institute of Public Health (NIPH) for generating high-quality genomic data. The research is part of the HARVEST collaboration, supported by the Research Council of Norway (#229624). We also thank the NORMENT Centre for providing genotype data, funded by the Research Council of Norway (#223273), South East Norway Health Authorities and Stiftelsen Kristian Gerhard Jebsen, and in collaboration with deCODE Genetics. We further thank the Center for Diabetes Research, the University of Bergen for providing genotype data funded by the ERC AdG project SELECTionPREDISPOSED, Stiftelsen Kristian Gerhard Jebsen, Trond Mohn Foundation, the Research Council of Norway, the Novo Nordisk Foundation, the University of Bergen, and the Western Norway Health Authorities. The MoBa work was performed on the TSD (Tjeneste for Sensitive Data) facilities, owned by the University of Oslo, operated and developed by the TSD service group at the University of Oslo, IT Department (USIT). EY is supported by the European Union (Grant numbers: 101045526, 101073237) and the Research Council of Norway (Grant numbers: 336078, 288083, 331640).

We would like to acknowledge the participants and investigators of the Generation Scotland Cohort study. Generation Scotland received core support from the Chief Scientist Office of the Scottish Government Health Directorates [CZD/16/6] and the Scottish Funding Council [HR03006]. Genotyping and methylation typing of the GS:SFHS samples was carried out by the Genetics Core Laboratory at the Wellcome Trust Clinical Research Facility, Edinburgh, Scotland and was funded by the Medical Research Council UK and the Wellcome Trust (Wellcome Trust Strategic Award “STratifying Resilience and Depression Longitudinally” (STRADL) Reference 104036/Z/14/Z).

We would like to thank and acknowledge the participants and investigators of the Estonian Biobank (EstBB) study. The research was conducted using the Estonian Center of Genomics/Roadmap II funded by the Estonian Research Council (project number TT17)

Norwegian analyses were performed on resources provided by Sigma2 - the National Infrastructure for High-Performance Computing and Data Storage in Norway. Estonian Data analysis was carried out in the High-Performance Computing Center cloud provided by University of Tartu. Analysis of the Generation Scotland data and the summary statistics obtained from the other analyses was conducted at IST Austria and is supported by the Scientific Service Units (SSU) of IST Austria through resources provided by Scientific Computing (SciComp).

## Declarations

## Author contributions

IK and MRR conceived and designed the study. IK developed the computer code for JODIE and the within-family GWAS, with input from MRR and MM. MRR conducted the Estonian biobank imputation and phasing, with oversight from RJH and OD. IK and MRR analyzed the Estonian and Generation Scotland cohort data, with oversight from AC, CH, and RM. IK conducted the Generation Scotland phasing. LH conducted the Norwegian mother, father and child study phasing, imputation and analysis with input from MRR, IK, ECC, and AH. Estonian data collection, genotyping and QC was conducted by the Estonian Biobank research team: AM, LM, TE, RM, MN and GH. AH provided study oversight and contributed the MoBa data. MRR, IK, LH and AH wrote the paper. All authors approved the final manuscript prior to submission.

## Author consortia

The Estonian Biobank research team consist of Andres Metspalu, Lili Milani, Tõnu Esko, Reedik Mägi, Mari Nelis and Georgi Hudjashov.

## Data availability

Information on how to access the MoBaPsychGen post-imputation QC data is available here: https://www.fhi.no/en/more/research-centres/psychgen/access-to-genetic-data-after-quality-control-by-the-mobapsychgen-pipeline-v/.

Estonian Biobank data (https://genomics.ut.ee/en/content/estonian-biobank) were used in this project. For access to be granted to the Estonian Biobank genotypic and corresponding phenotypic data, a preliminary application must be presented to the oversight committee, who must first approve the project. Ethics permission must then be obtained from the Estonian Committee on Bioethics and Human Research. Finally, a full project must be submitted and approved by the Estonian Biobank.

Access to the Generation Scotland data is available with appropriate permission from the Generation Scotland Access Committee. Applications should be made to. (https://www.ed.ac.uk/lothian-birth-cohorts/data-access-collaboration).

Haplotype Reference Consortium Release 1.1 data (https://ega-archive.org/datasets/EGAD00001002729) are available by application to a Data Access Committee (DAC) of the Wellcome Trust Sanger Institute.

The Common Metabolic Diseases Atlas can be accessed here https://cmdga.org.

## Code availability

Source code for the phasing, imputation, and analysis is available at https://github.com/medical-genomics-group/jodie for the JODIE model. plink version 1.9 is available at https://www.cog-genomics.org/plink/. KING is available at https://www.kingrelatedness.com. SHAPEIT5 is available at https://github.com/odelaneau/shapeit5.

## Supplementary Information

**Fig. S1.**
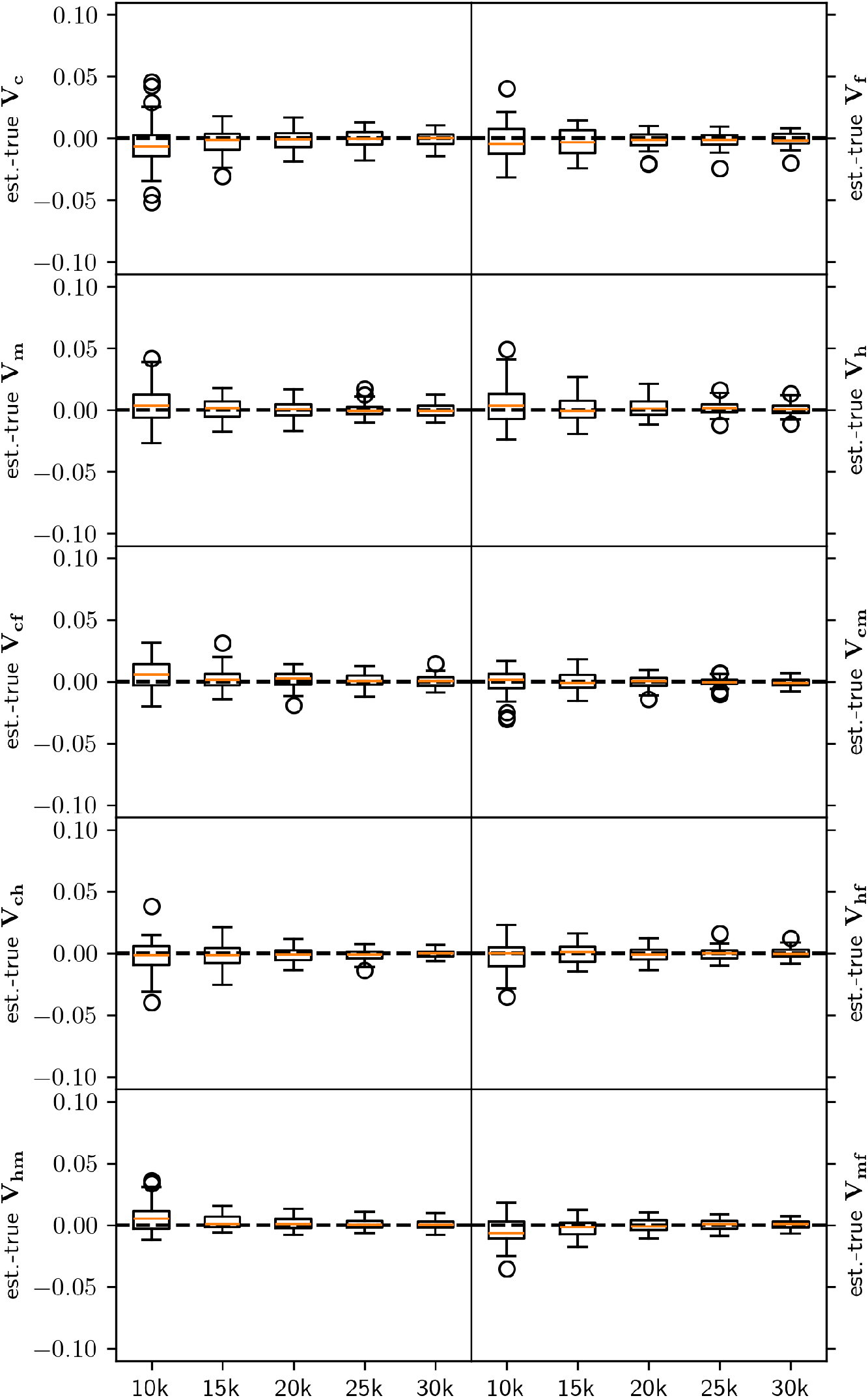
Box plot of difference between estimated and truly simulated variances for different sample sizes. The difference between estimated and true variances and covariances for the realistic variance scenario, *V*_5_, are shown for 50 simulation replicates under random mating for n=10k, 15k, 20k, 25k, 30k individuals and p=52,310 SNPs selected from the 1000 human genome project.

**Fig. S2.**
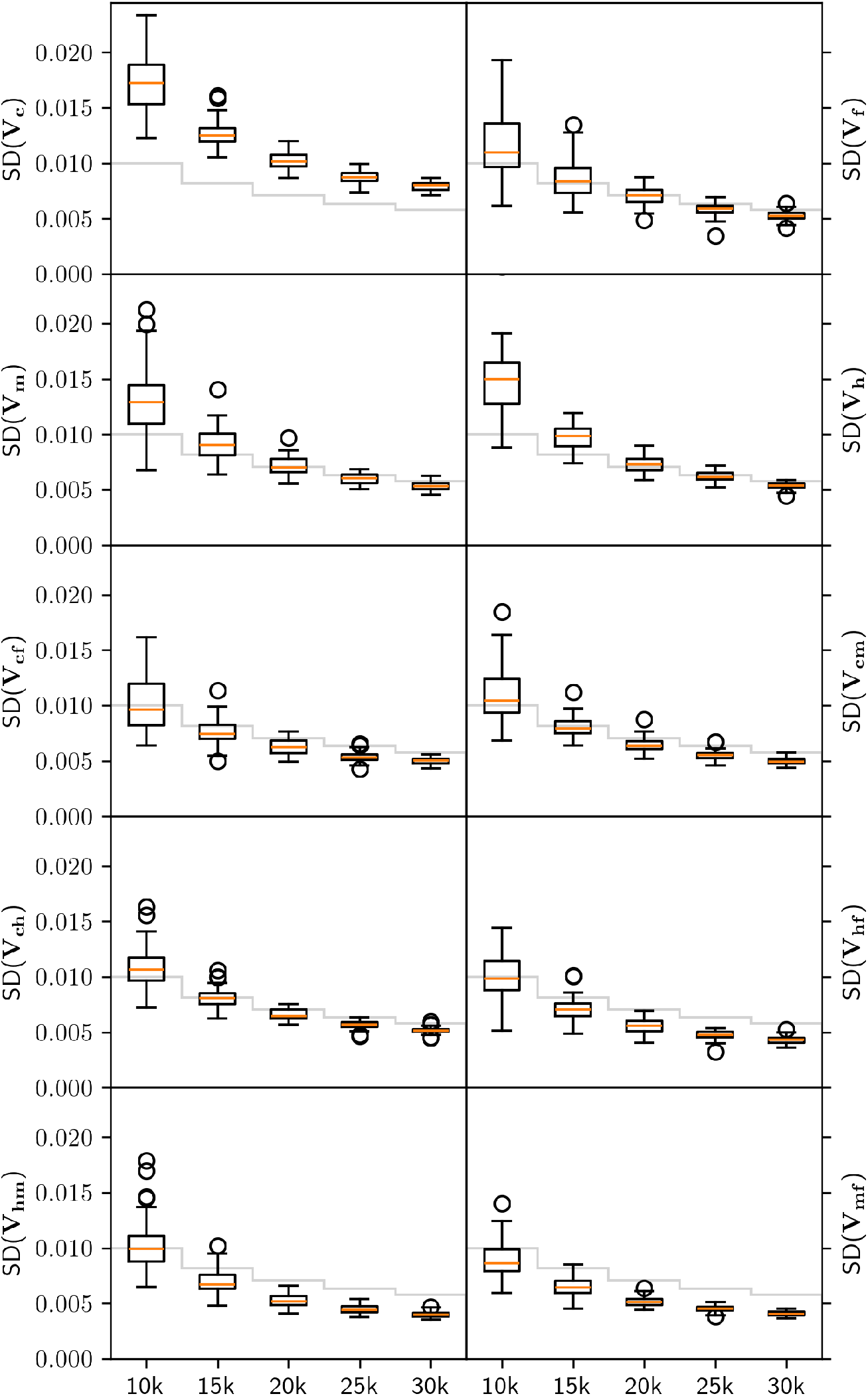
Box plot of estimated standard deviation of variances in simulation study for different sample sizes. The standard deviation of the estimated variances and covariances across posterior iterations for the realistic variance scenario, *V*_5_, are shown for 50 simulation replicates under random mating for n=10k, 15k, 20k, 25k, 30k individuals and p=52,310 SNPs selected from the 1000 human genome project. The grey lines indicate the Poissonian error 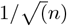, directly reflecting the uncertainty due to the sample size. Note that the estimated standard deviation is the square root of variation in posterior estimates across posterior iterations, thus only indirectly reflects the sample size.

**Fig. S3.**
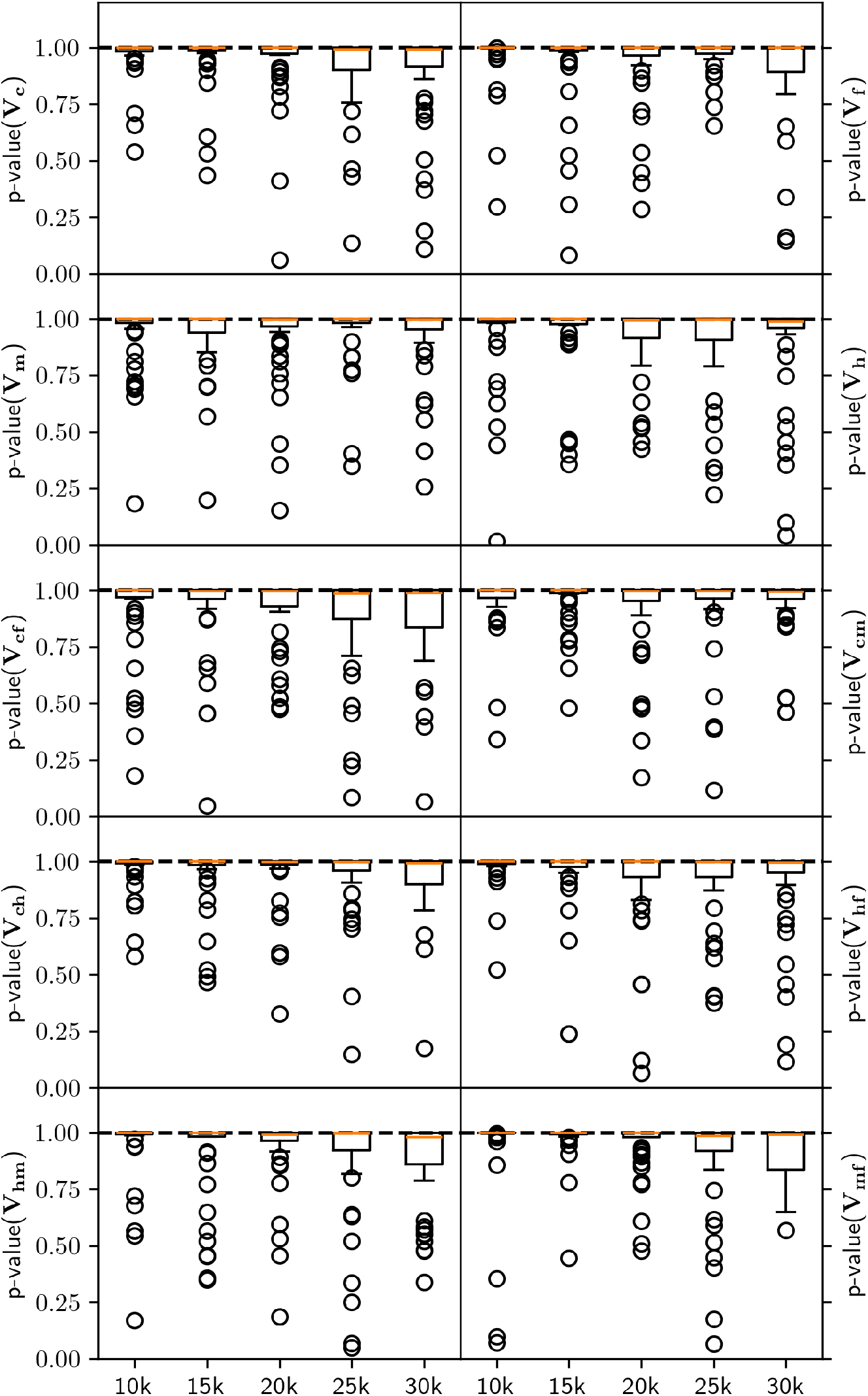
Box plot of p-value for simulation based calibration test for different sample sizes. To assess the soundness of our sampler, we run a simulation based calibration test for different sample sizes and the realistic variance scenario, *V*_5_. For each of the 50 simulation replicates, we create histograms of rank statistics that conform to a uniform distribution if the sampler is well calibrated. We test the uniformity of ranks with a *χ*^2^ test. The p-values of this test are shown for n=10k, 15k, 20k, 25k, 30k individuals and p=52,310 SNPs selected from the 1000 human genome project.

**Fig. S4.**
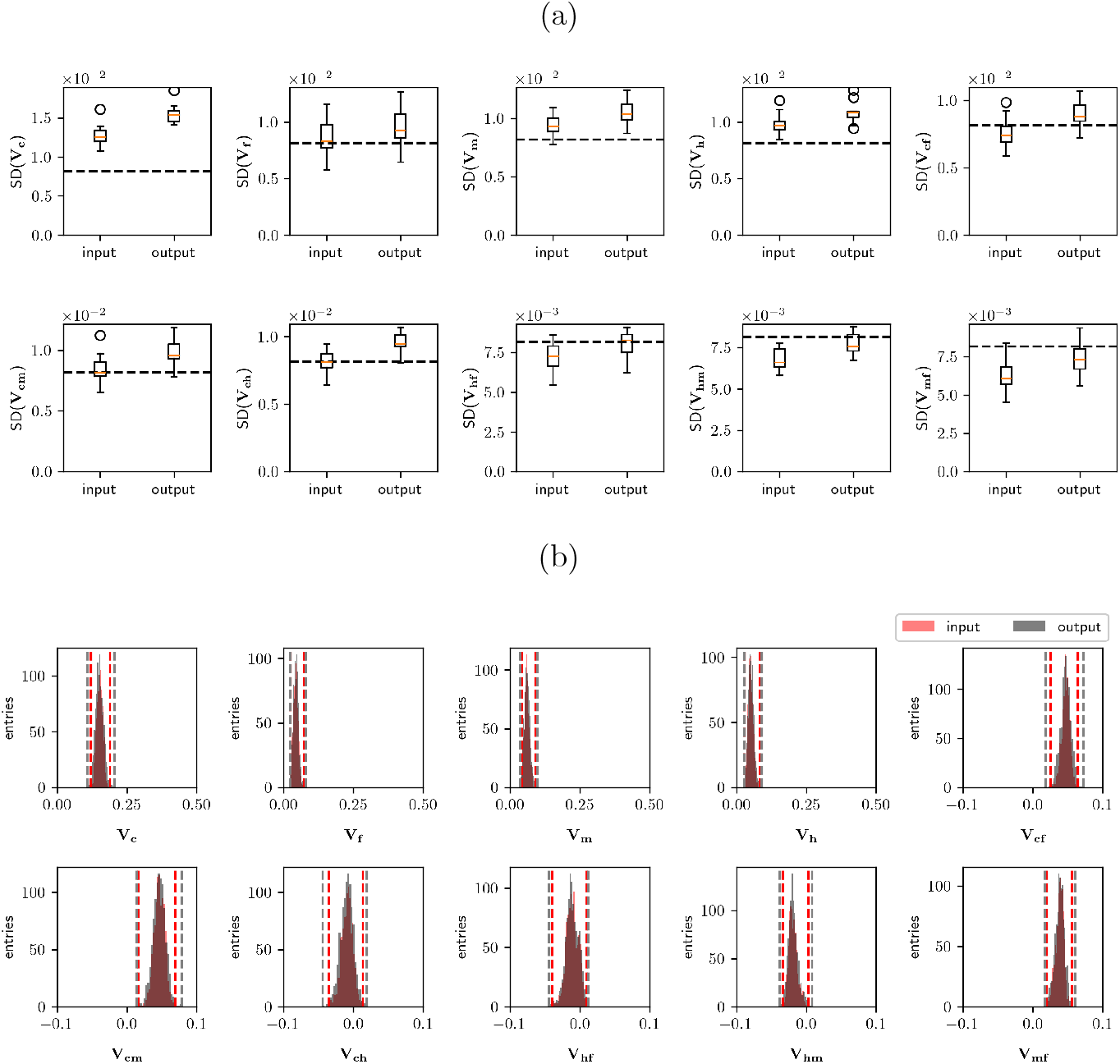
Example of posterior predictive checks. For the realistic V scenario, ***V***_**5**_, n=15,000 individuals and p=52,310 SNPs selected from the 1000 human genome project, the variance elements of all 1000 posterior iterations estimated with JODIE (”input”) are used to regenerate the effects and resample all elements of ***V*** (”output”) with the same function used in JODIE. The standard deviations (SD) of all variance elements across 10 simulation replicates of each variance element is shown in (a). The dashed lines indicate the Poissonian error, 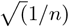. For one of the 10 simulations, the input and output distributions of all iterations are shown in (b). The grey dashed lines indicate the minimum and maximum values of the output which always encompasses the minimum and maximum values of the input, shown as red dashed lines.

**Fig. S5.**
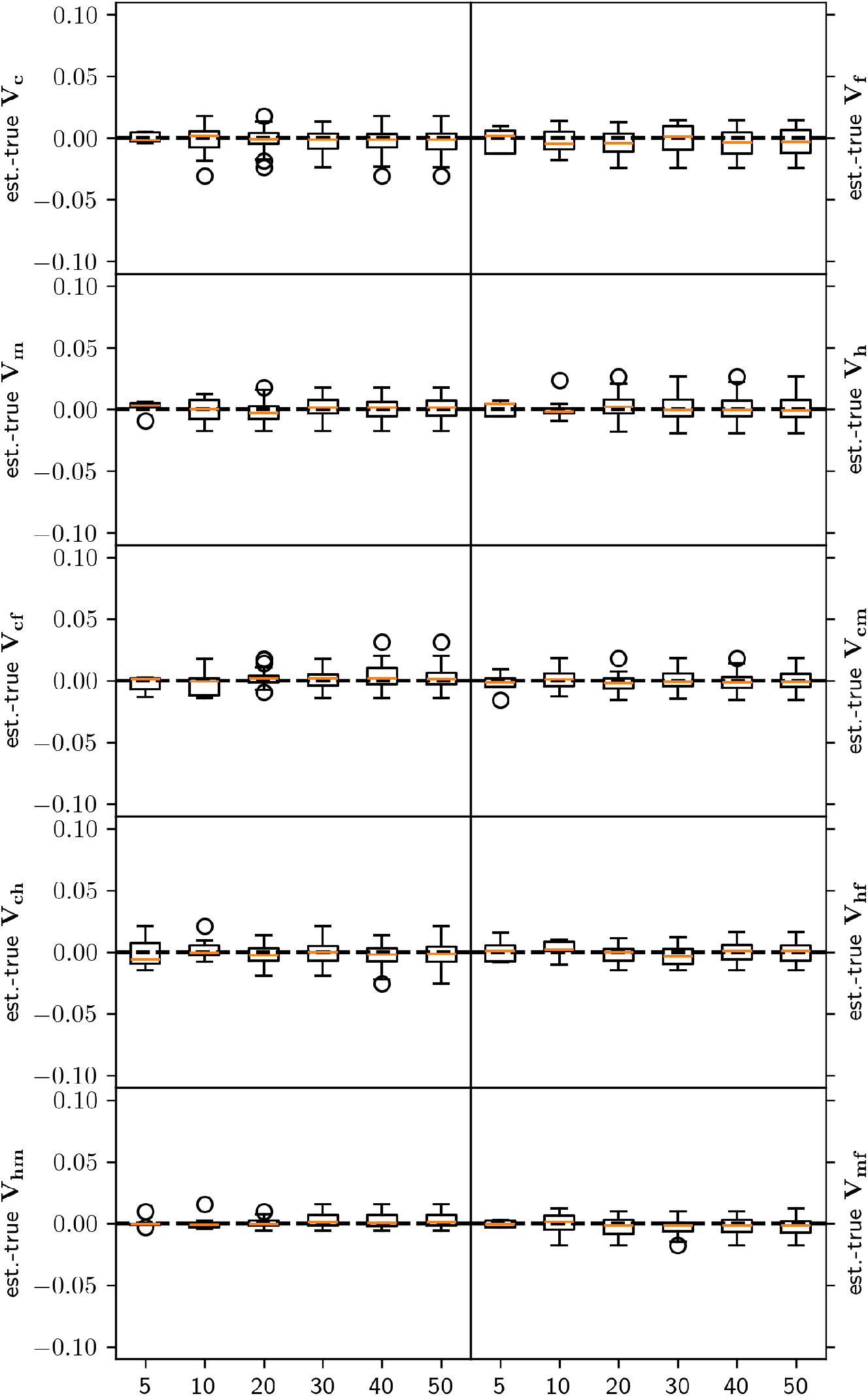
Box plot of difference between estimated and truly simulated variances for different number of simulation replicates. The difference between estimated and true variances and covariances for the realistic variance scenario, ***V***_**5**_, are shown for 5, 10, 20, 30, 40 and 50 simulation replicates under random mating for n=15,000 individuals and p=52,310 SNPs selected from the 1000 human genome project. This shows that the number of simulation replicates does not bias the variance estimates. We chose to use 10 simulation replicates, as running simulations are computationally expensive.

**Fig. S6.**
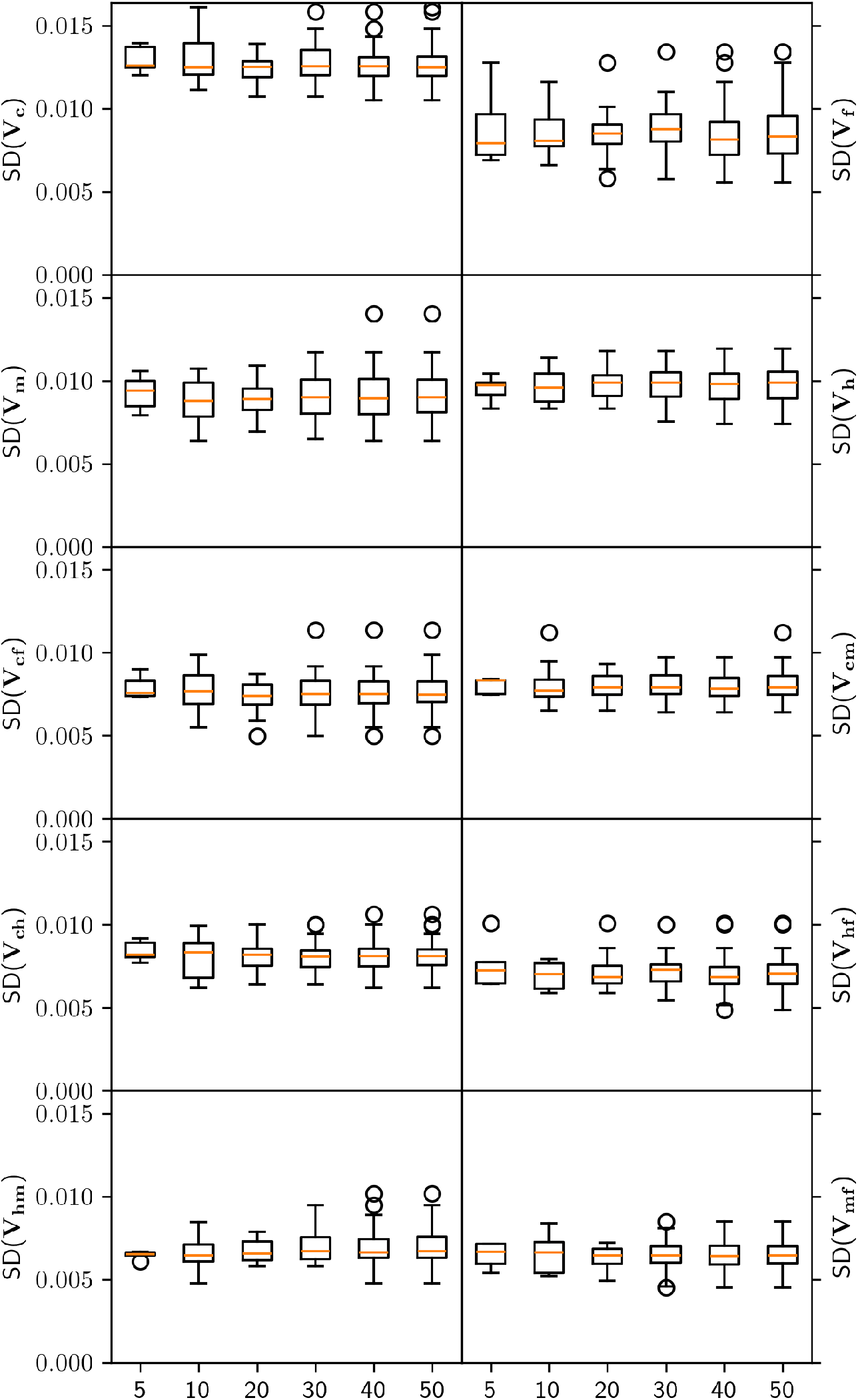
Box plot of estimated standard deviation of variances in simulation study for different number of simulation replicates. The standard deviation of the estimated variances and covariances across posterior iterations for the realistic variance scenario, ***V***_**5**_, are shown for 5, 10, 20, 30, 40 and 50 simulation replicates under random mating for n=15,000 individuals and p=52,310 SNPs selected from the 1000 human genome project. This shows that the standard deviation of variance estimates does not differ significantly across different numbers of simulation replicates. We chose to use 10 simulation replicates, as running simulations are computationally expensive.

**Fig. S7.**
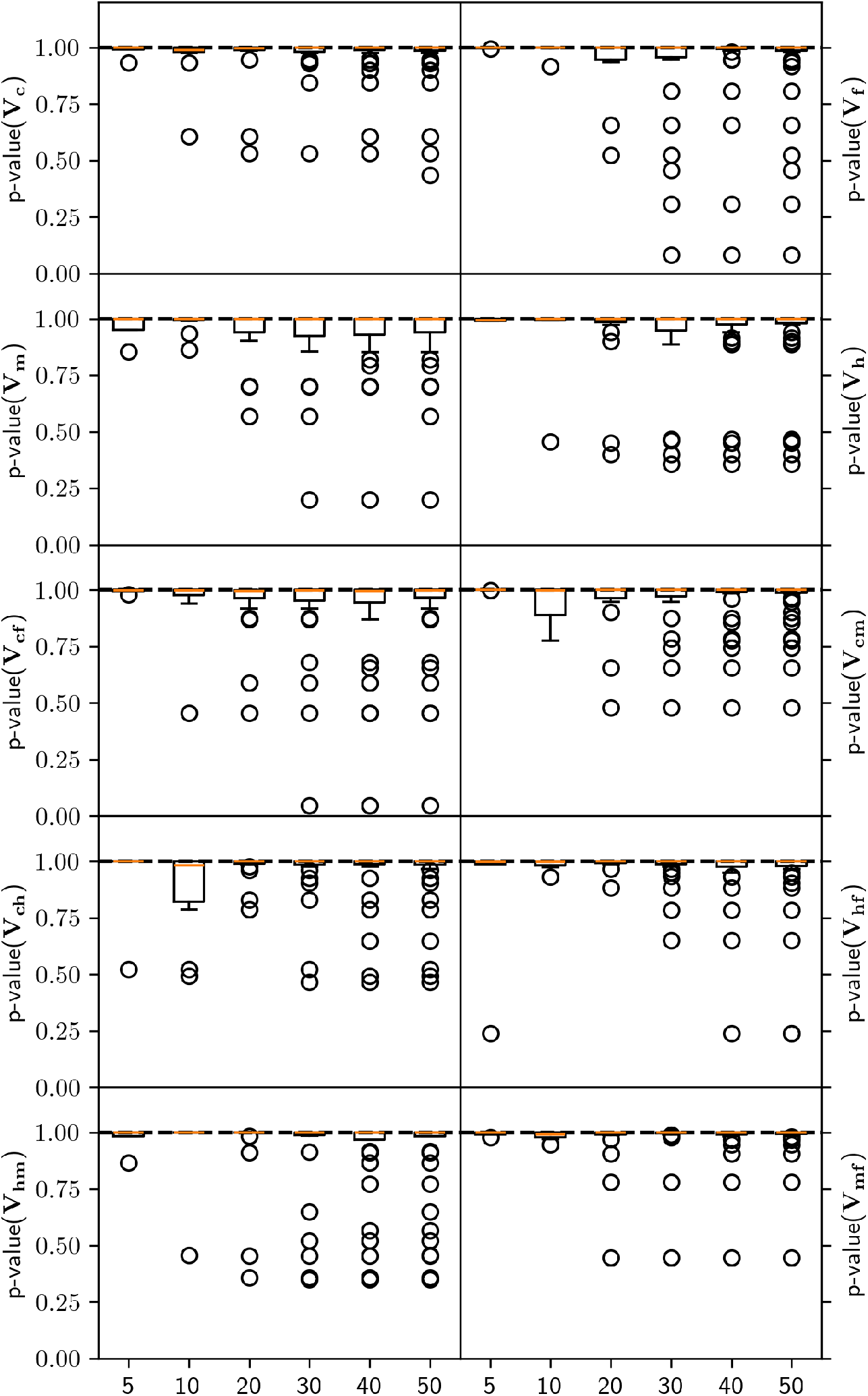
Box plot of p-value for simulation based calibration test for different numbers of simulation replicates. To assess the soundness of our sampler, we run a simulation based calibration test for different number of simulation replicates and the realistic variance scenario, ***V***_**5**_. For each simulation replicate, we create histograms of rank statistics that conform to a uniform distribution if the sampler is well calibrated. We test the uniformity of our ranks with a *χ*^2^ test. The p-values of this test are shown for n=15,000 individuals, p=52,310 SNPs selected from the 1000 human genome project and 5, 10, 20, 30, 40 and 50 simulation replicates, randomly drawn from the total 50 replicates. The sampler is always well calibrated, independently of number of simulation replicates.

**Fig. S8.**
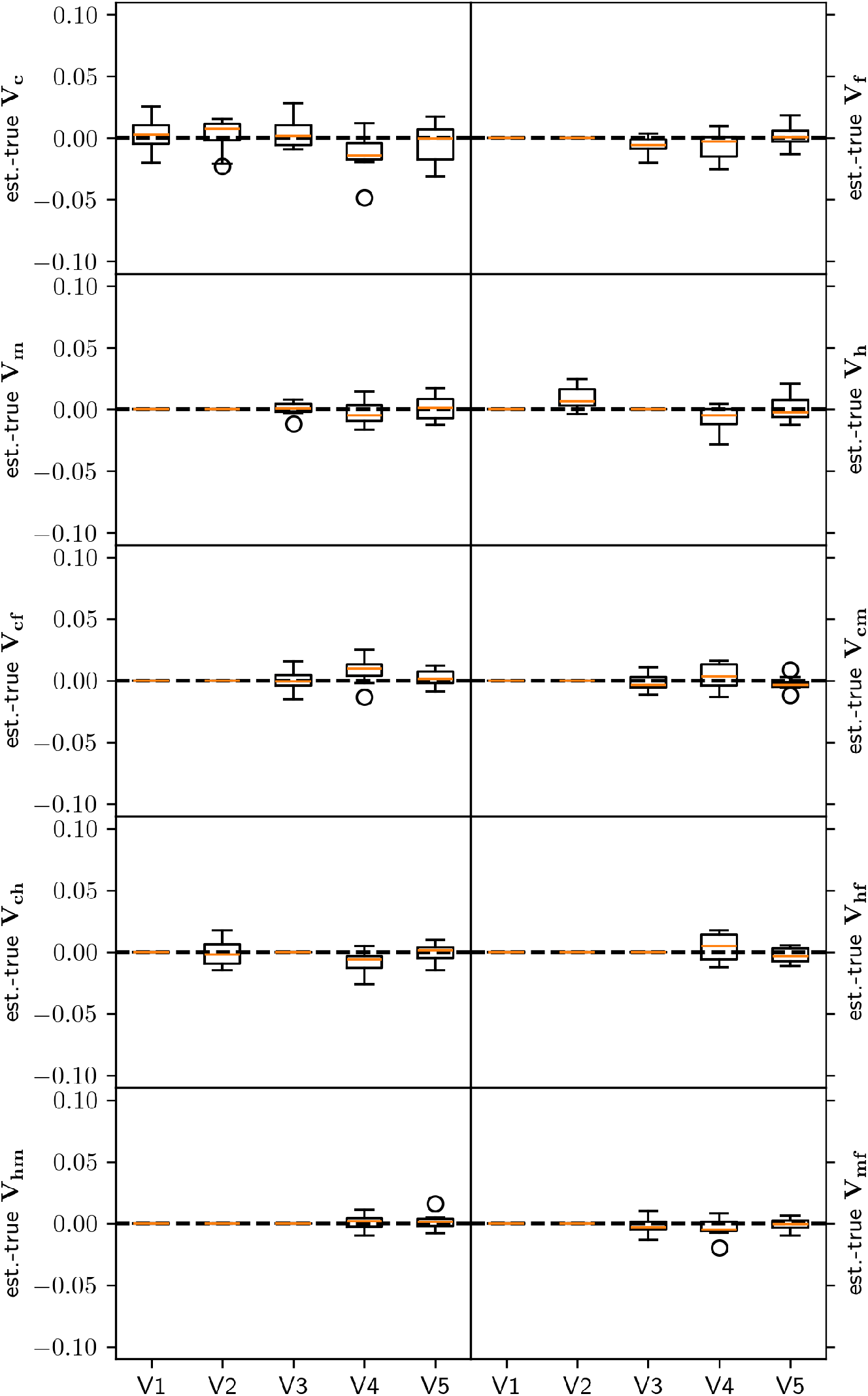
Box plot of difference between estimated and truly simulated variances, applying the region of practical equivalence (ROPE) rule. For the five different simulation scenarios, the difference between estimated and true variances and covariances are shown for 10 simulation replicates under random mating for n=16,000 individuals and p=52,310 SNPs selected from the 1000 human genome project. A significance threshold on the variances (and corresponding covariances) of 2% for more than 95% of the iterations was applied.

**Fig. S9.**
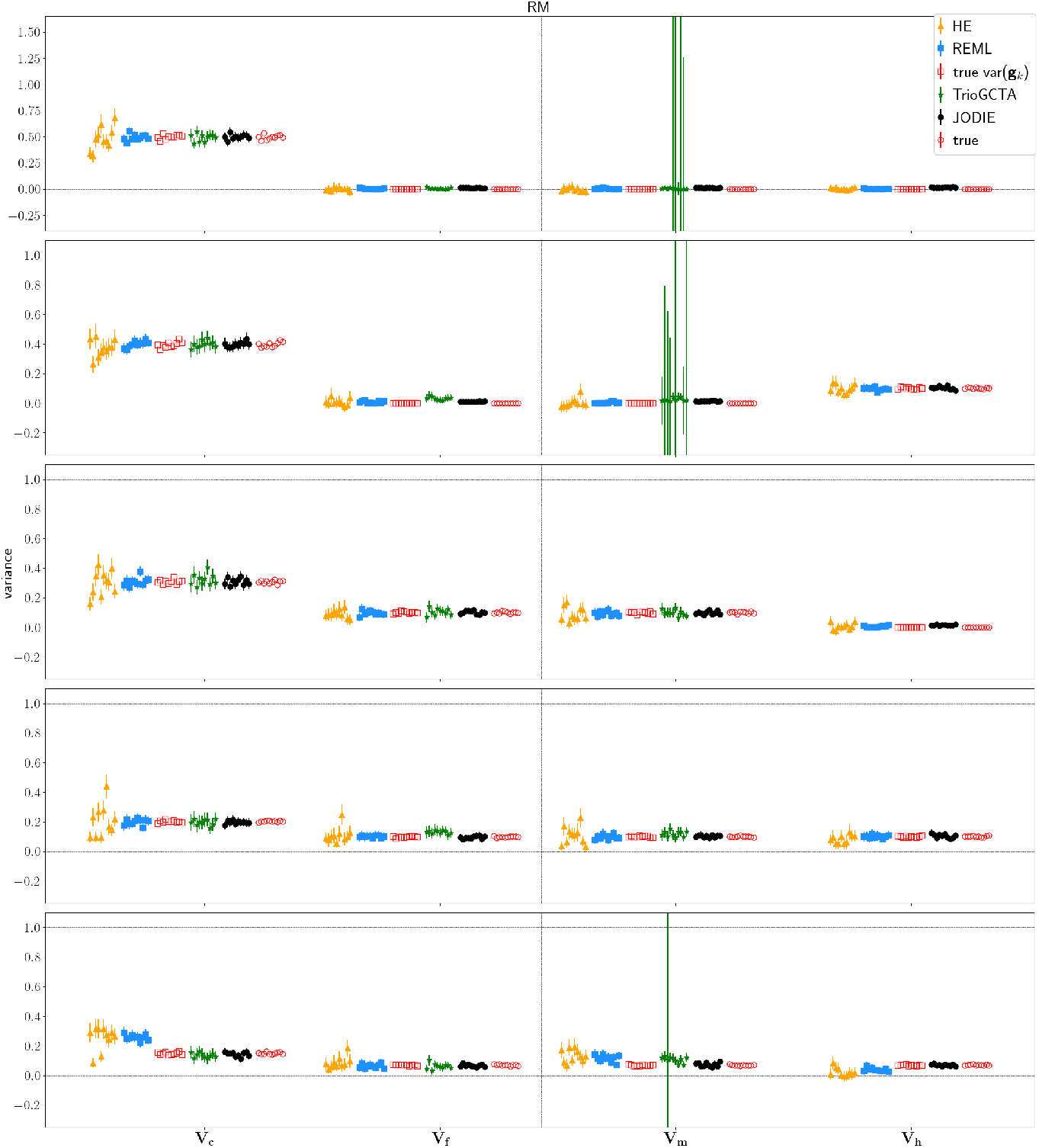
Simulation study of the influence of random mating on the estimation of variances using various statistical methods. For five different simulation scenarios (top to bottom), true variances of effects, true variance of ***Xβ*** for each genetic component (labeled ”true var(**g**_*k*_)”) and variances estimated with Haseman-Elston regression (HE), REML, TrioGCTA and JODIE for 10 simulations under random mating are shown for n=16,000 individuals and p=52,310 SNPs selected from the 1000 human genome project. Error bars represent 2× standard deviation and 95% confidence intervals, respectively.

**Fig. S10.**
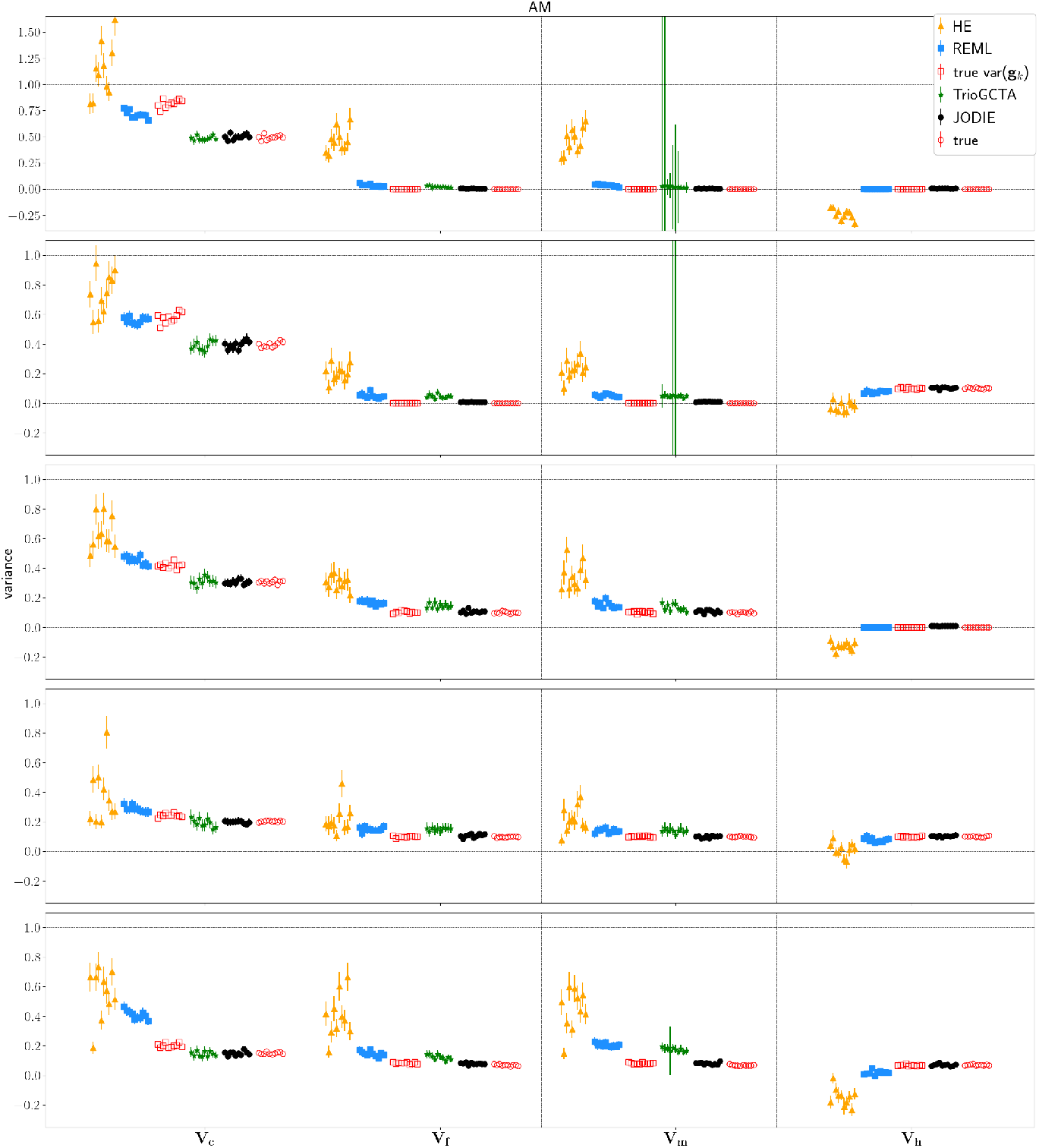
Simulation study of the influence of assortative mating on the estimation of variances using various statistical methods. For five different simulation scenarios (top to bottom), true variances of effects, true variance of ***Xβ*** for each genetic component (labeled ”true var(**g**_*k*_)”) and variances estimated with Haseman-Elston regression (HE), REML, TrioGCTA and JODIE for 10 simulations under assortative mating are shown for n=16,000 individuals and p=52,310 SNPs selected from the 1000 human genome project. Error bars represent 2× standard deviation and 95% confidence intervals, respectively.

**Fig. S11.**
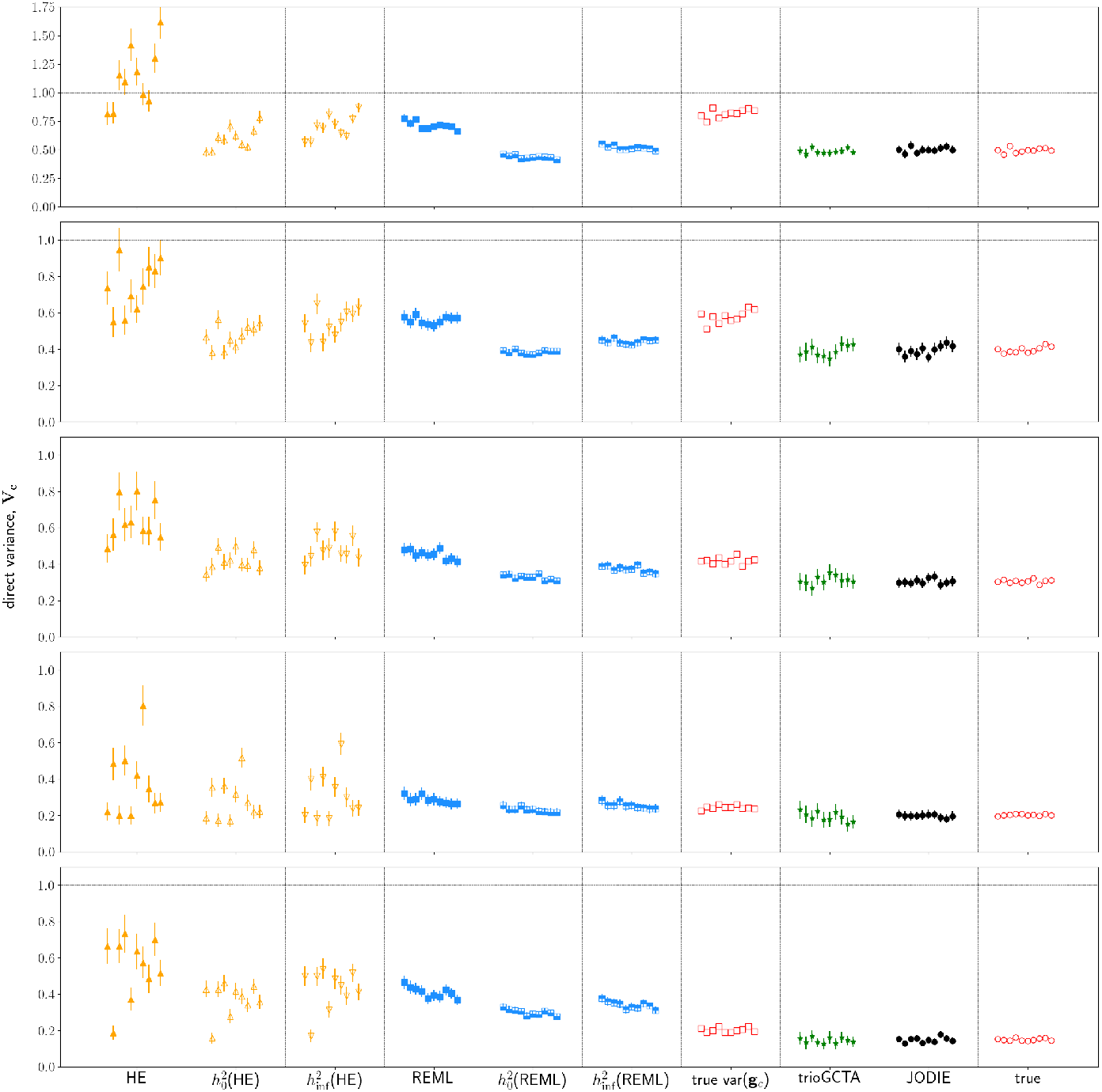
Simulation study of the influence of assortative mating on the estimation of the variance attributable to direct effects using various statistical methods, including bias correction. For five different simulation scenarios (top to bottom), variances attributable to direct effects estimated with Haseman-Elston regression (HE), REML, TrioGCTA, and JODIE, true variance of ***X***_***c***_***β***_***c***_ (labeled ”true var(**g**_*c*_)”), and true variances of direct effects (labelled ”true”), for 10 simulations under assortative mating are shown for n=16,000 individuals and p=52,310 SNPs selected from the 1000 human genome project. Results from HE and REML corrected for AM according to Ref. [20] (labeled 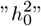 and 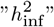) are also shown. Error bars represent 2× standard deviation and 95% confidence intervals, respectively.

**Fig. S12.**
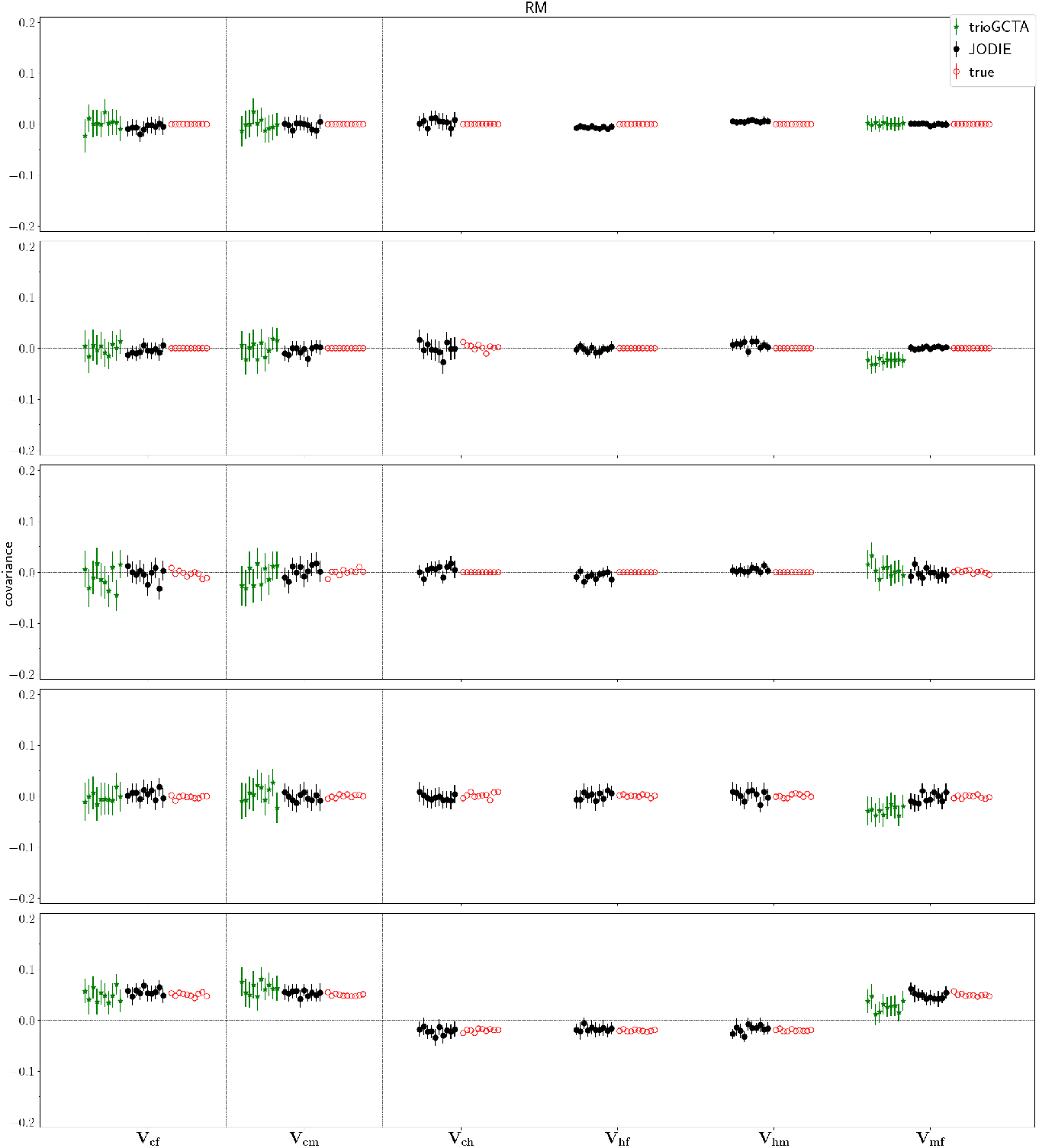
Simulation study of the estimation of covariances using JODIE. For five different simulation scenarios (top to bottom), the true and estimated covariances obtained with JODIE and TrioGCTA for 10 simulation replicates under random mating are shown for n=16,000 individuals and p=52,310 SNPs selected from the 1000 human genome project. Error bars represent 95% confidence intervals.

**Fig. S13.**
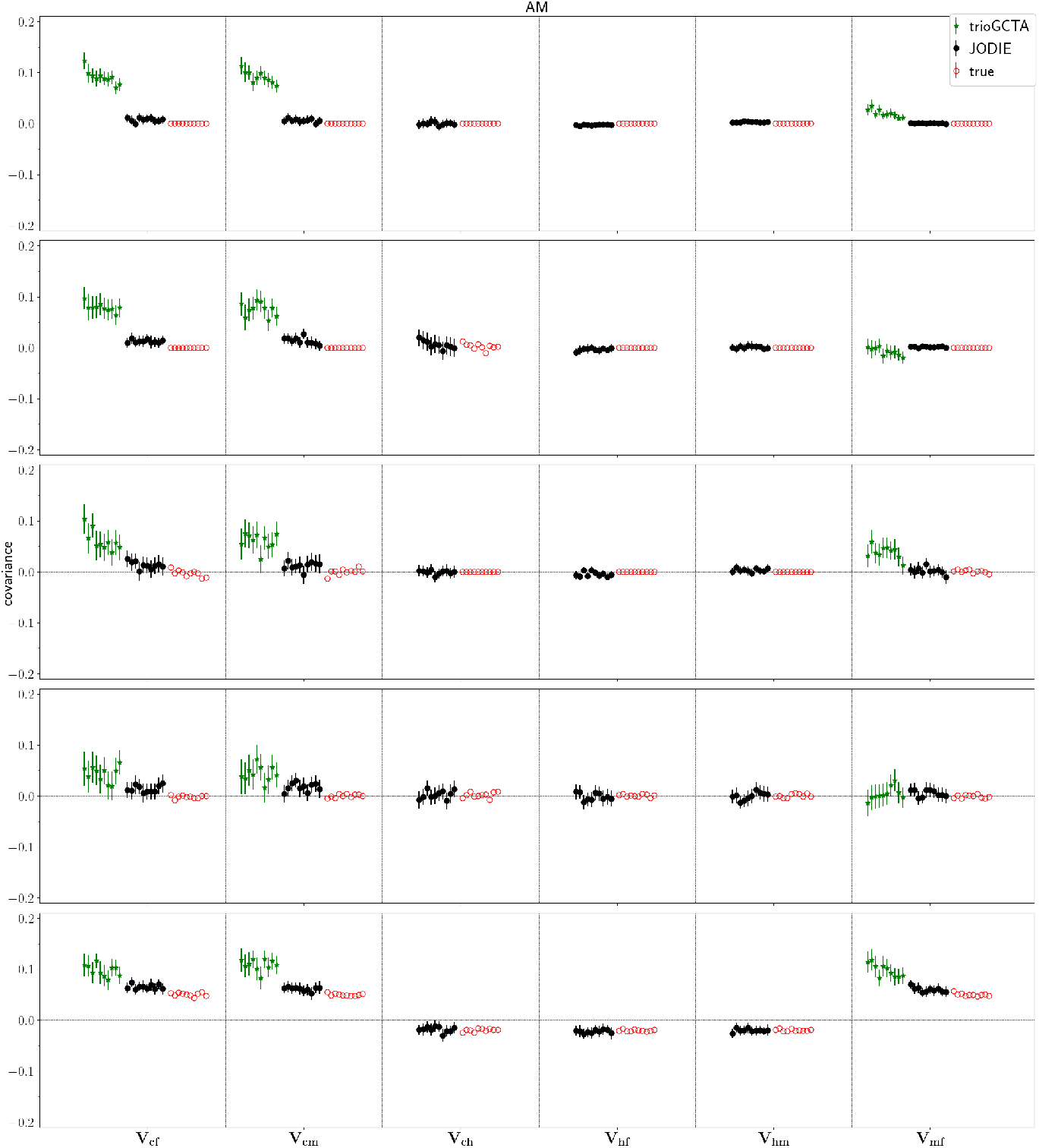
Simulation study of the influence of assortative mating on the estimation of covariances using JODIE and TrioGCTA. For five different simulation scenarios (top to bottom), the true and estimated covariances obtained with JODIE and TrioGCTA for 10 simulation replicates under assortative mating are shown for n=16,000 individuals and p=52,310 SNPs selected from the 1000 human genome project. Error bars represent 95% confidence intervals.

**Fig. S14.**
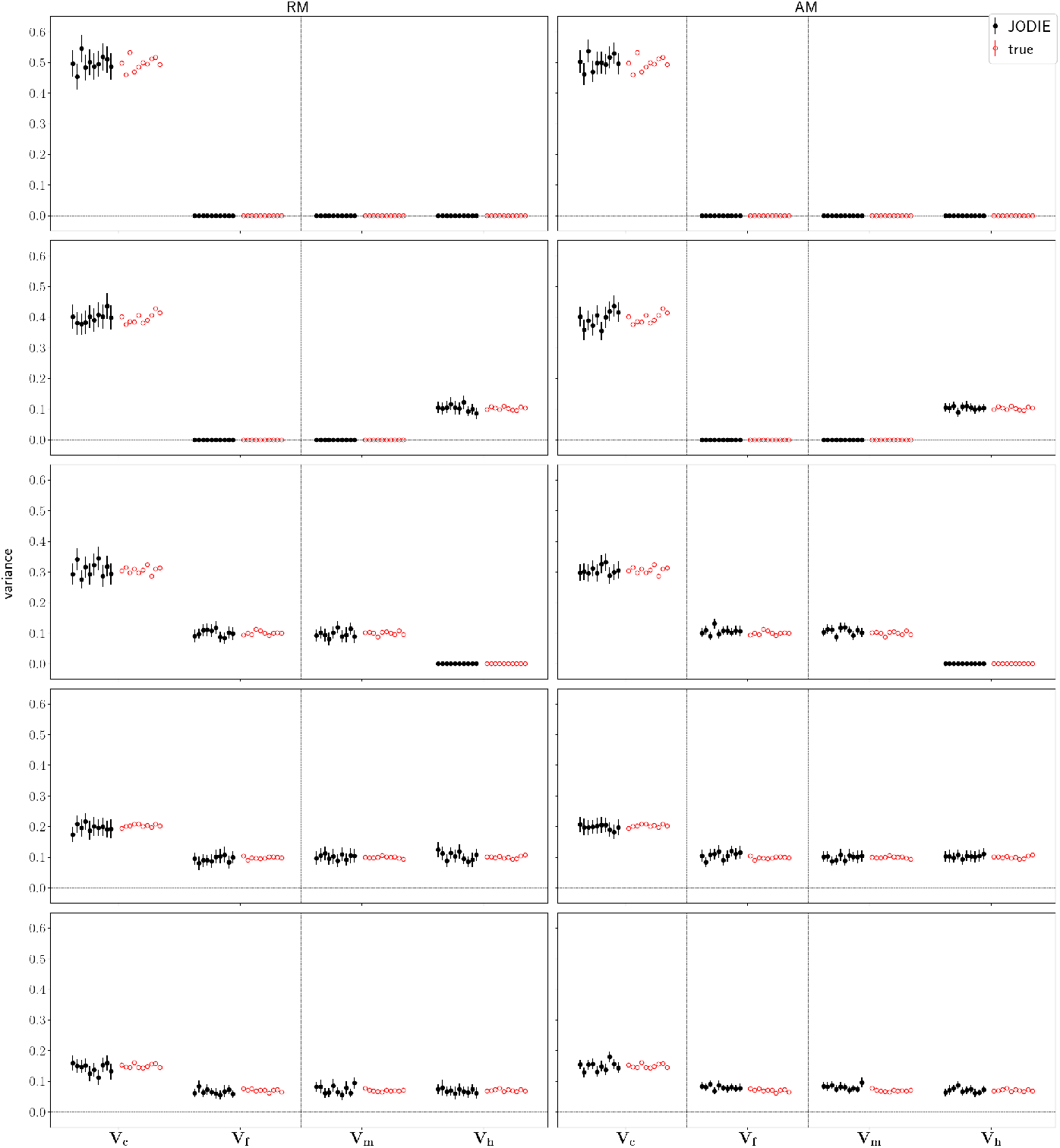
Simulation study for the estimation of variances applying the region of practical equivalence (ROPE) rule. For five different simulation scenarios (top to bottom), true and estimated variances obtained with JODIE for 10 simulation replicates under random (left) and assortative mating (right) are shown for n=16,000 individuals and p=52,310 SNPs selected from the 1000 human genome project. A significance threshold on the variances of 2% for more than 95% of the iterations was applied. Error bars represent 95% confidence intervals.

**Fig. S15.**
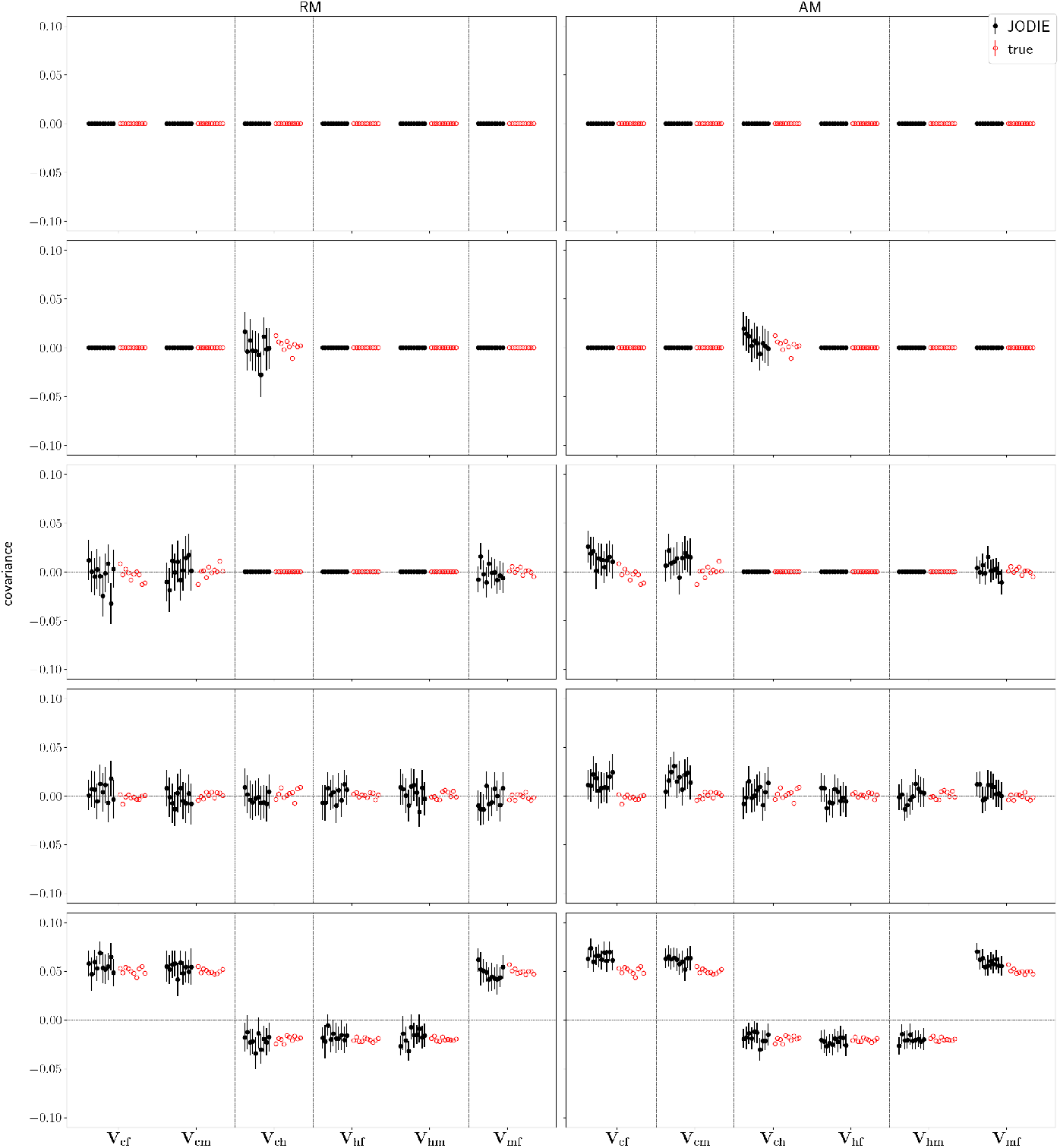
Simulation study for the estimation of covariances applying the region of practical equivalence (ROPE) rule. For five different simulation scenarios (top to bottom), true and estimated covariances obtained with JODIE for 10 simulation replicates are shown for n=16,000 individuals and p=52,310 SNPs selected from the 1000 human genome project. A significance threshold on the variances (and corresponding covariances) of 2% for more than 95% of the iterations was applied. Error bars represent 95% confidence intervals.

**Fig. S16.**
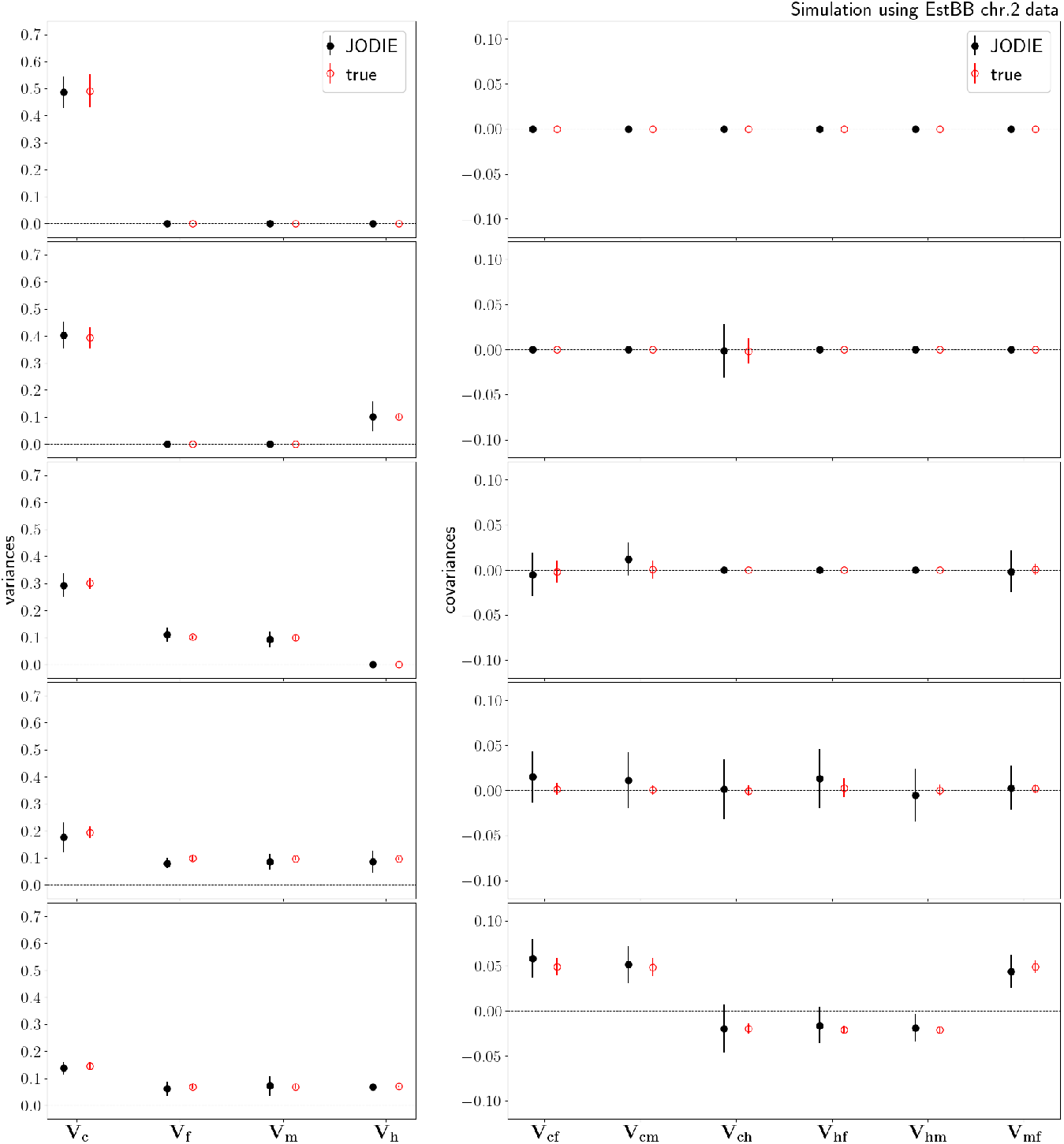
Real-data simulation study applying the region of practical equivalence (ROPE) rule. For the five different simulation scenarios (top to bottom), the average true and estimated variances (left) and covariances (right), obtained with JODIE, are shown for 10 simulation replicates using EstBB chromosome 2 with p=89,578 SNPs and n=10,512 individuals. A significance threshold on the variances (and corresponding covariances) of 2% for more than 95% of the iterations was applied. Error bars represent 95% confidence intervals.

**Fig. S17.**
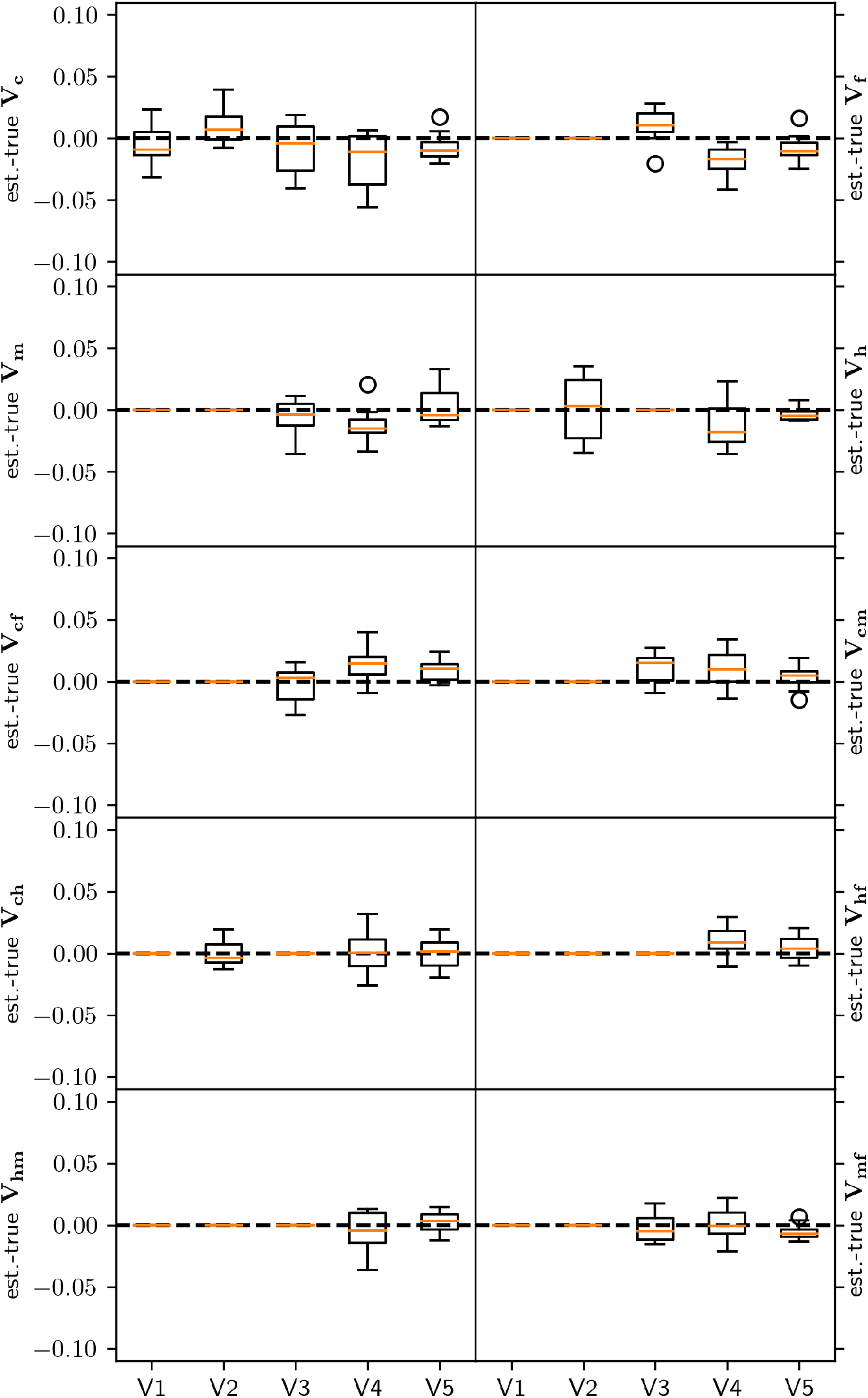
Box plot of real-data simulation study applying the region of practical equivalence (ROPE) rule. For the five different simulation scenarios, the difference between estimated and true variances and covariances are shown for 10 simulation replicates using EstBB chromosome 2 with p=89,578 SNPs and n=10,512 individuals. A significance threshold on the variances (and corresponding covariances) of 2% for more than 95% of the iterations was applied.

**Fig. S18.**
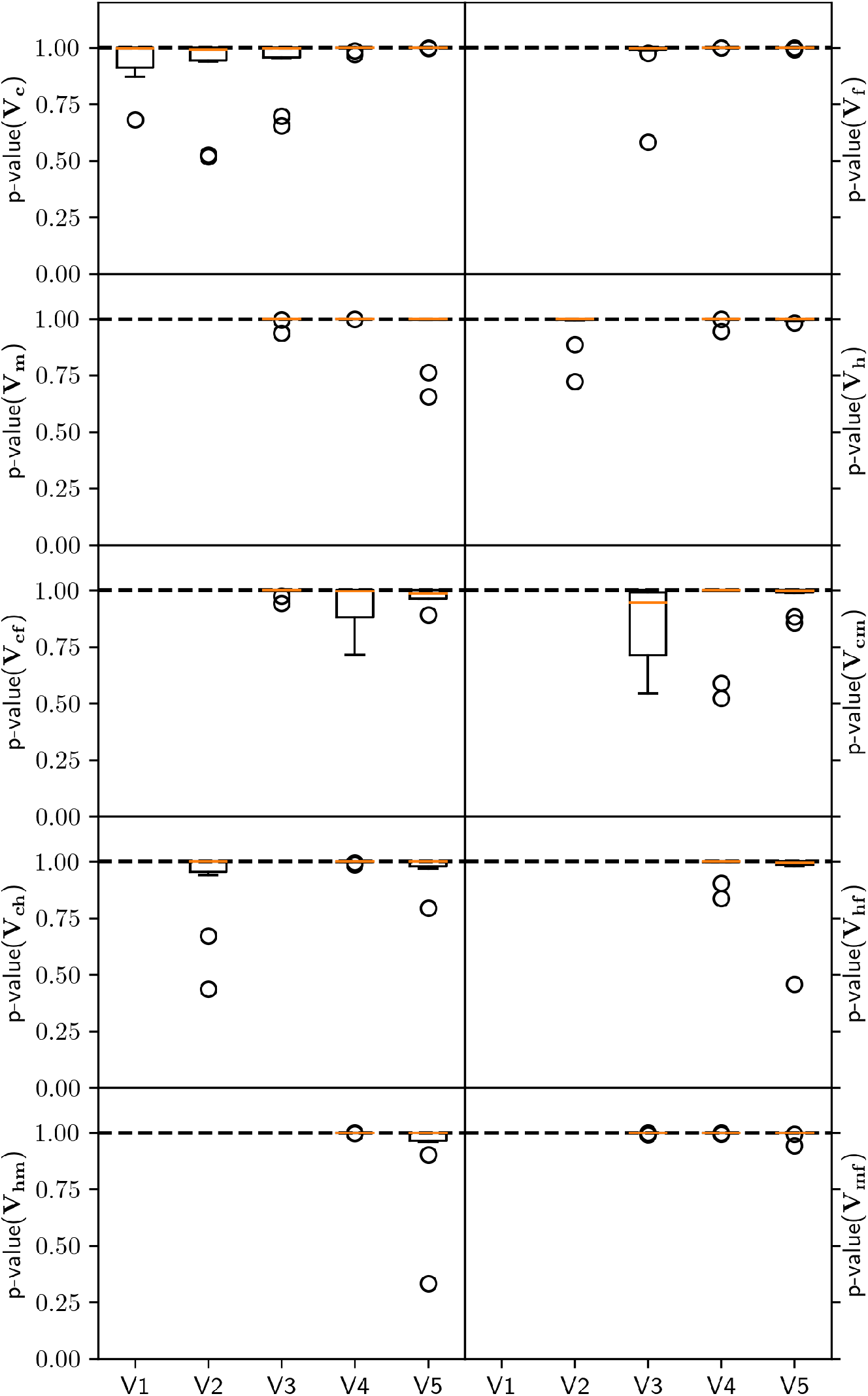
Box plot of p-value for simulation based calibration test for realdata simulations. To assess the soundness of our sampler, we run a simulation based calibration test for different variance scenarios, ***V***_**1**_ to ***V***_**5**_, using EstBB chromosome 2 with p=89,578 SNPs and n=10,512 individuals. For each simulation replicate, we create histograms of rank statistics that conform to a uniform distribution if the sampler is well calibrated. We test the uniformity of our ranks with a *χ*^2^ test. The p-values of this test show that the sampler is well calibrated under all variance scenarios.

**Fig. S19.**
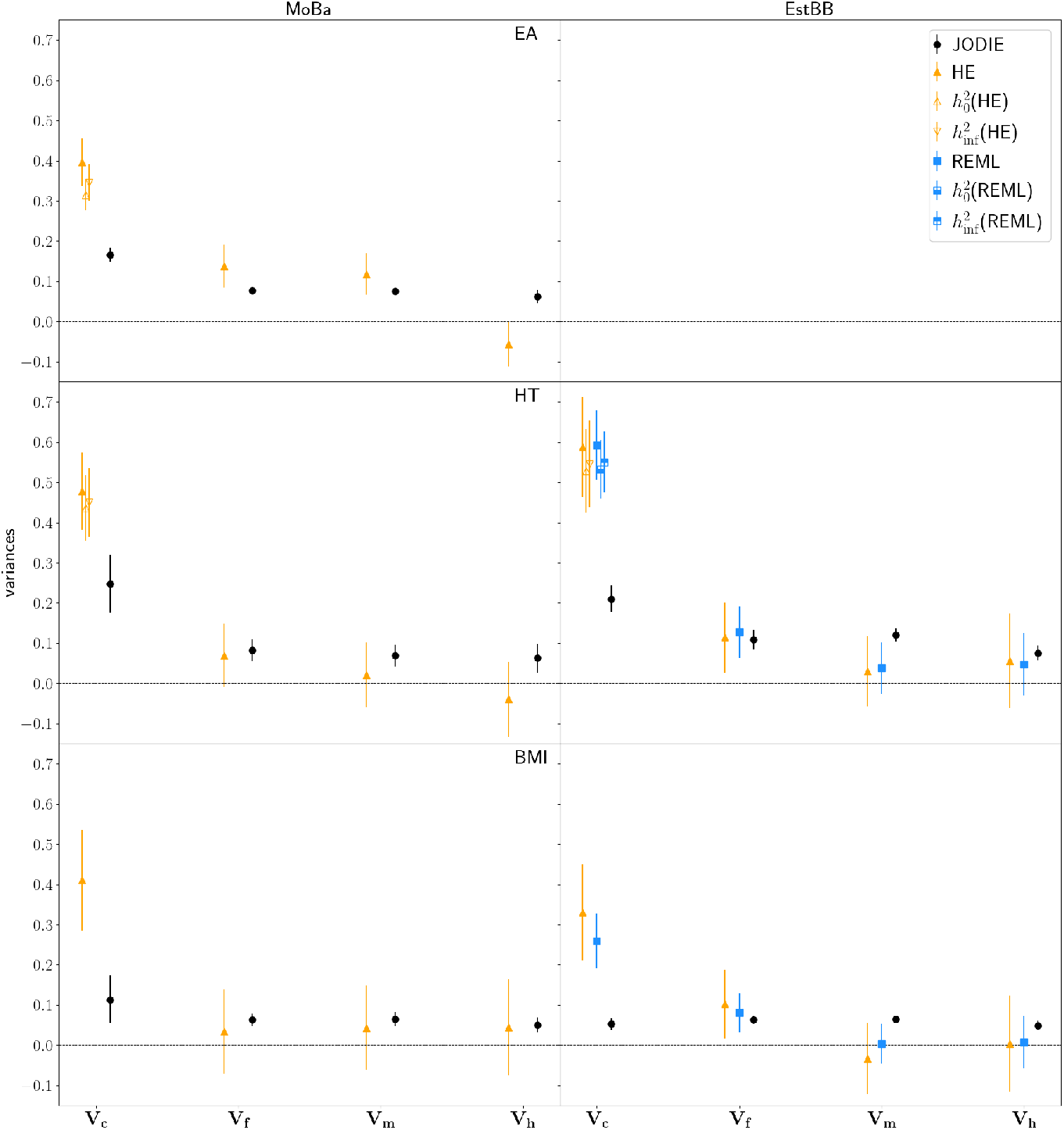
Comparison of genetic variances estimated with various statistical methods in MoBa and EstBB. Variances estimated with Haseman-Elston regression (HE), REML and JODIE in MoBa (left) and EstBB (right) for educational scores (EA, top), height (HT, middle) and body mass index (BMI, bottom). For HE and REML results for EA and HT, *V*_*c*_ is also shown corrected for AM according to Ref. [20] (labeled 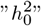 and 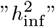) using genetic similarities between partners determined in MoBa [21]. Error bars represent 2*×*standard deviation and 95% confidence intervals, respectively. Note that the definition of variance differs depending on the method.

**Fig. S20.**
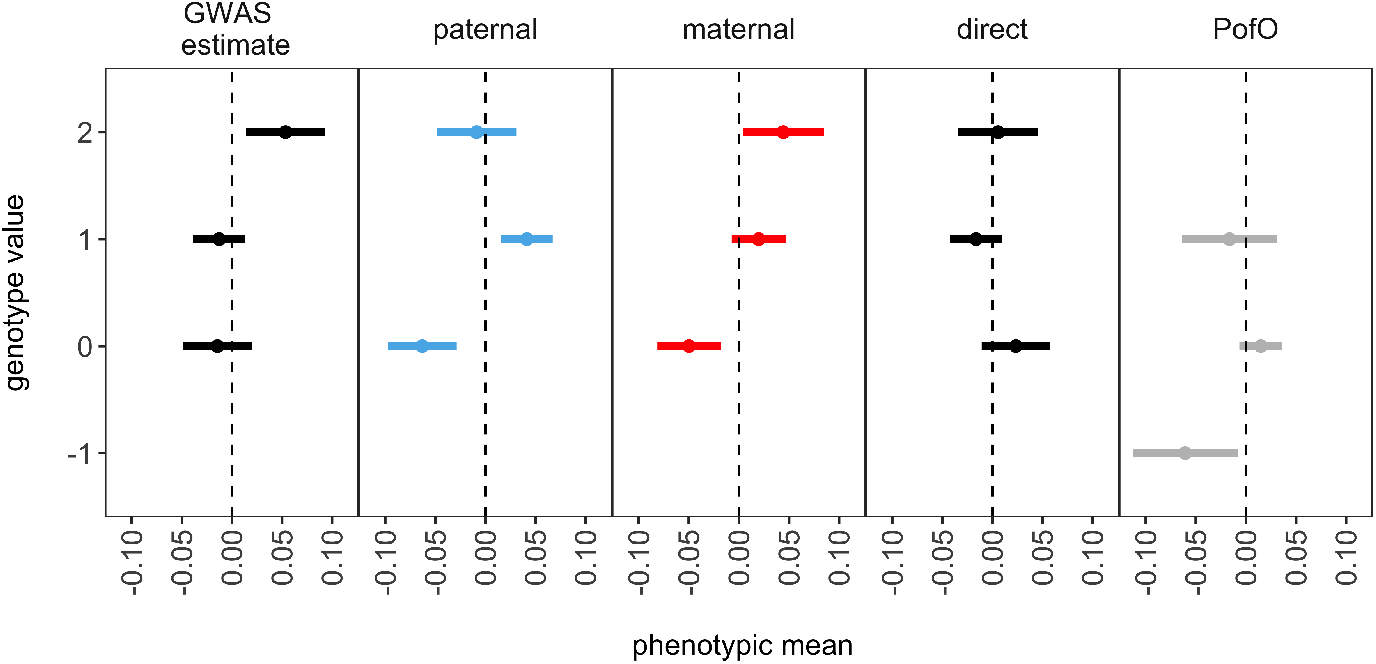
Mean years of schooling in the Generation Scotland (GS) study for individuals of different genotypes at rs13237829. In GS, we calculate the mean phenotypic value for years of education, standardized to zero mean and unit variance, for different child and parental genotypes at rs13237829. This SNP was identified as being significantly associated with education scores in an fGWAS analysis using MoBa data, and is significantly associated in the UK Biobank in a standard GWAS analysis (p-value 1.65e-06), and an fGWAS analysis of 2980 families in GS (p-value = 0.004). From left to right, the plots show the phenotypic means of children for (a) children of different genotypic values (0,1,2) from direct effects (labeled ”GWAS estimate”), reflecting the regression coefficient estimates from a simple marginal ordinal least squares regression of the phenotype onto the genotypic values of the child; (b) different parental genotypic effects (labeled ”paternal” and ”maternal”); (c) children of different genotypic values conditional on the parental genotype effects (labeled ”direct”); (d) heterozygous children who inherit the minor allele from their fathers (value of 1) or from their mothers (value of -1), with all other genotypes scored as 0. Error bars represent the standard error of the mean.

**Fig. S21.**
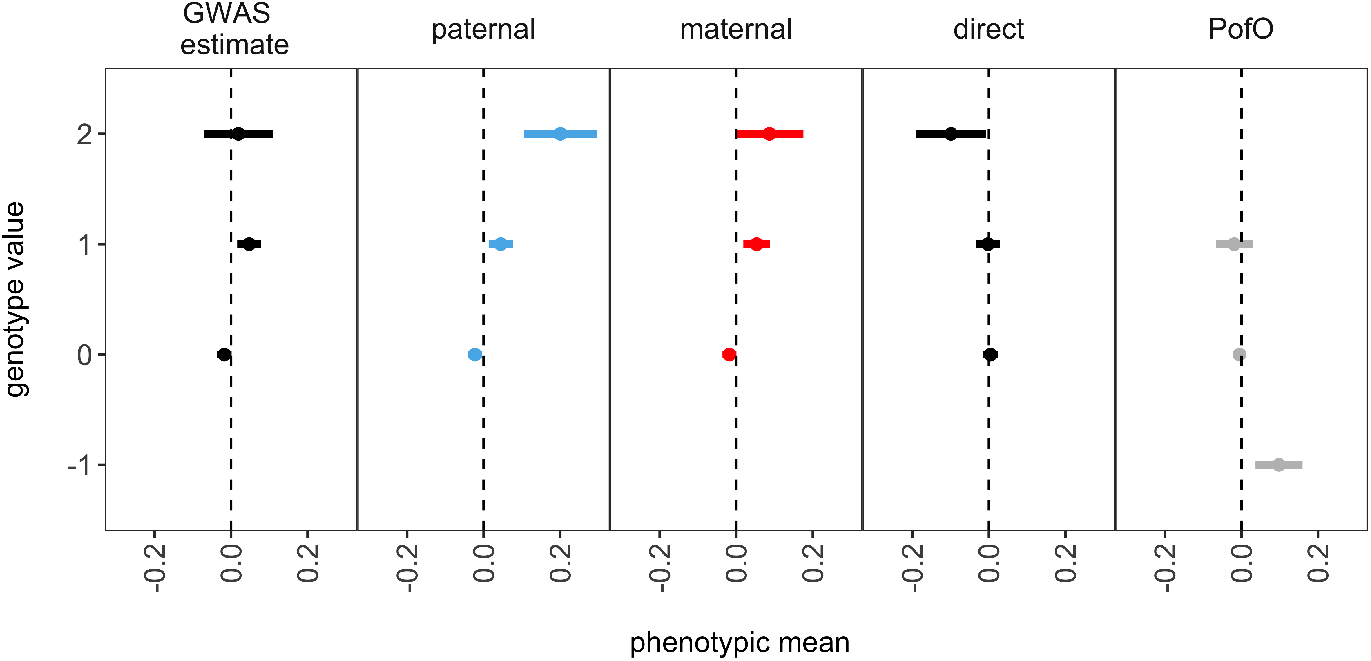
Mean sex- and age-adjusted standing height (HT) in the Generation Scotland (GS) study for individuals of different genotypes at rs646806. In GS, we calculate the mean phenotypic value for standing height, adjusted for age and sex and standardized to zero mean and unit variance, for different child and parental genotypes at rs646806. This SNP was identified as being significantly associated with height in an fGWAS analysis using MoBa data (p-value 1.11e-08), and is significantly associated in the UK Biobank in a standard GWAS analysis (p-value 1.49e-39), and an fGWAS analysis of 2980 families in GS (p-value = 0.001). From left to right, the plots show the phenotypic means of children for (a) children of different genotypic values (0,1,2) from direct effects (labeled ”GWAS estimate”), reflecting the regression coefficient estimates from a simple marginal ordinal least squares regression of the phenotype onto the genotypic values of the child; (b) different parental genotypic effects (labeled ”paternal” and ”maternal”); (c) children of different genotypic values conditional on the parental genotype effects (labeled ”direct”); (d) heterozygous children who inherit the minor allele from their fathers (value of 1) or from their mothers (value of -1), with all other genotypes scored as 0. Error bars represent the standard error of the mean.

**Fig. S22.**
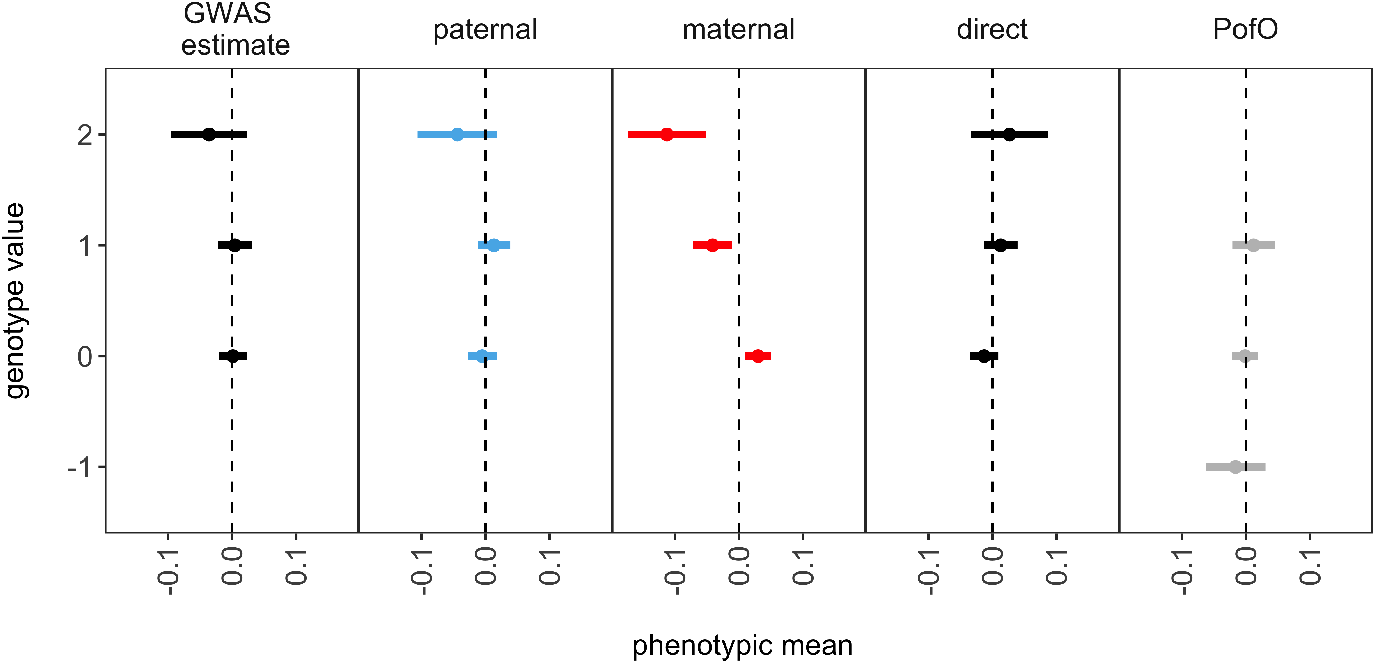
Mean sexand age-adjusted body mass index (BMI) in the Generation Scotland (GS) study for individuals of different genotypes at rs919918. In GS, we calculate the mean phenotypic value for BMI, adjusted for age and sex and standardized to zero mean and unit variance, for different child and parental genotypes at rs919918. This SNP was identified as being significantly associated with height in an fGWAS analysis using MoBa data (p-value 2.14e-08), and is significantly associated in the UK Biobank in a standard GWAS analysis (p-value 4.29e-06), and an fGWAS analysis of 2980 families in GS (p-value = 0.003). From left to right, the plots show the phenotypic means of children for (a) children of different genotypic values (0,1,2) from direct effects (labeled ”GWAS estimate”), reflecting the regression coefficient estimates from a simple marginal ordinal least squares regression of the phenotype onto the genotypic values of the child; (b) different parental genotypic effects (labeled ”paternal” and ”maternal”); (c) children of different genotypic values conditional on the parental genotype effects (labeled ”direct”); (d) heterozygous children who inherit the minor allele from their fathers (value of 1) or from their mothers (value of -1), with all other genotypes scored as 0. Error bars represent the standard error of the mean.

## Supplementary Tables

**Table S1:**
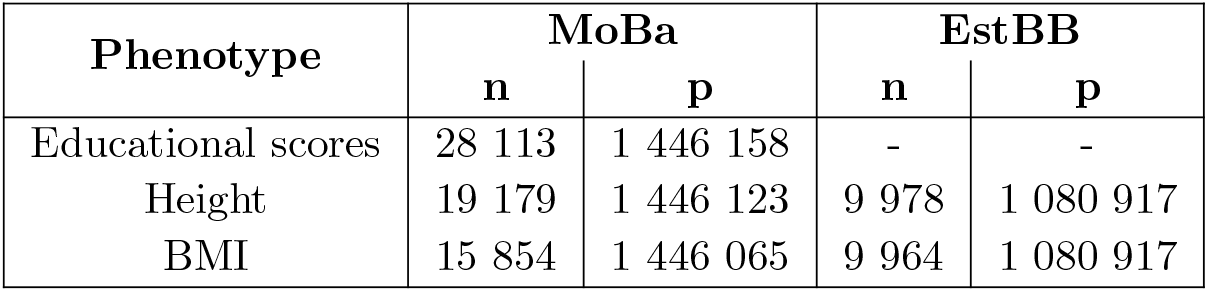
Overview of datasets. Number of individuals, n, and number of markers, p, for each phenotype in MoBa and EstBB.

**Table S2:** Estimated covariance values from JODIE. Estimated posterior mean genetic (co)variances ± uncertainty (95% credible interval) for educational scores (EA), height (HT) and body mass index (BMI) in MoBa and EstBB. Covariances where the mean ± uncertainty include the value 0 are not shown.

| (co)v. | EA | HT |  | BMI |  |
| --- | --- | --- | --- | --- | --- |
|  | MoBa | MoBa | EstBB | MoBa | EstBB |
| $\hat{V}_c$ | $0.17 \pm 0.02$ | $0.25 \pm 0.07$ | $0.21 \pm 0.03$ | $0.11 \pm 0.06$ | $0.05 \pm 0.01$ |
| $\hat{V}_f$ | $0.08 \pm 0.01$ | $0.08 \pm 0.03$ | $0.11 \pm 0.02$ | $0.06 \pm 0.02$ | $0.06 \pm 0.01$ |
| $\hat{V}_m$ | $0.08 \pm 0.01$ | $0.07 \pm 0.03$ | $0.12 \pm 0.02$ | $0.06 \pm 0.02$ | $0.06 \pm 0.01$ |
| $\hat{V}_h$ | $0.06 \pm 0.02$ | $0.06 \pm 0.04$ | $0.08 \pm 0.02$ | $0.05 \pm 0.02$ | $0.05 \pm 0.01$ |
| $\hat{V}_{cf}$ | $0.06 \pm 0.01$ | $0.05 \pm 0.02$ | $0.06 \pm 0.02$ | — | — |
| $\hat{V}_{cm}$ | $0.06 \pm 0.01$ | — | $0.07 \pm 0.01$ | — | — |
| $\hat{V}_{mh}$ | — | — | $-0.03 \pm 0.01$ | — | — |
| $\hat{V}_{fm}$ | $0.03 \pm 0.02$ | — | — | — | — |

**Table S3:**
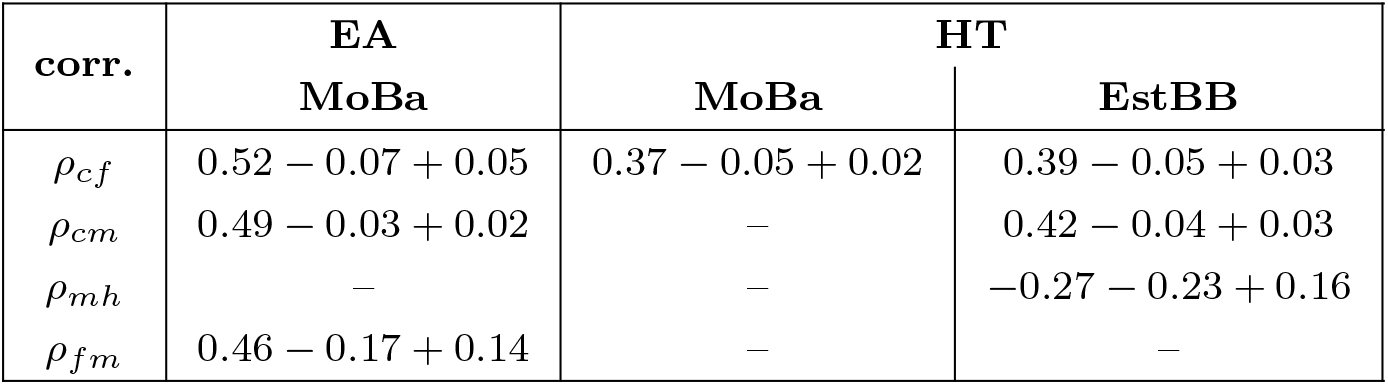
Correlation values from JODIE. Genetic correlations calculated from the estimated posterior mean genetic covariances and their lower and upper uncertainties for educational scores (EA) and height (HT) in MoBa and EstBB. Correlations where the mean genetic covariance ± uncertainty include the value 0, are not shown, which is the case all correlation for BMI. The upper and lower uncertainties are calculated using the 95% credible intervals instead of the means.

| **corr.** | **EA** | **HT** |  |
| --- | --- | --- | --- |
|  | **MoBa** | **MoBa** | **EstBB** |
| $\rho_{cf}$ | $0.52 - 0.07 + 0.05$ | $0.37 - 0.05 + 0.02$ | $0.39 - 0.05 + 0.03$ |
| $\rho_{cm}$ | $0.49 - 0.03 + 0.02$ | – | $0.42 - 0.04 + 0.03$ |
| $\rho_{mh}$ | – | – | $-0.27 - 0.23 + 0.16$ |
| $\rho_{fm}$ | $0.46 - 0.17 + 0.14$ | – | – |

